# Cellular signaling beyond the Wiener-Kolmogorov limit

**DOI:** 10.1101/2021.07.15.452575

**Authors:** Casey Weisenberger, David Hathcock, Michael Hinczewski

## Abstract

Accurate propagation of signals through stochastic biochemical networks involves significant expenditure of cellular resources. The same is true for regulatory mechanisms that suppress fluctuations in biomolecular populations. Wiener-Kolmogorov (WK) optimal noise filter theory, originally developed for engineering problems, has recently emerged as a valuable tool to estimate the maximum performance achievable in such biological systems for a given metabolic cost. However, WK theory has one assumption that potentially limits its applicability: it relies on a linear, continuum description of the reaction dynamics. Despite this, up to now no explicit test of the theory in nonlinear signaling systems with discrete molecular populations has ever seen performance beyond the WK bound. Here we report the first direct evidence the bound being broken. To accomplish this, we develop a theoretical framework for multi-level signaling cascades, including the possibility of feedback interactions between input and output. In the absence of feedback, we introduce an analytical approach that allows us to calculate exact moments of the stationary distribution for a nonlinear system. With feedback, we rely on numerical solutions of the system’s master equation. The results show WK violations in two common network motifs: a two-level signaling cascade and a negative feedback loop. However the magnitude of the violation is biologically negligible, particularly in the parameter regime where signaling is most effective. The results demonstrate that while WK theory does not provide strict bounds, its predictions for performance limits are excellent approximations, even for nonlinear systems.

## 1 Introduction

Fundamental mathematical limits on the behavior of biochemical reaction networks^1–6^ provide fascinating insights into the design space of living systems. Though these limits remain notoriously permeable compared to their analogues in physics—subject to re-interpretaton and exceptions as additional biological complexities are discovered—they still give a rough guide to what is achievable by natural selection for a given set of resources. They also raise other interesting issues^7, 8^: is selection actually strong enough to push a particular system toward optimality? When is performance sacrificed due to metabolic costs or the randomizing forces of genetic drift?

Information processing in cellular networks has been a particularly fertile ground for discussing optimality. Certain cellular processes like environmental sensing rely on accurate information transfer through intrinsically stochastic networks of reactions^9, 10^. Other processes in development and regulation depend on suppressing noise through homeostatic mechanisms like negative feedback^11–14^. Either scenario, whether maintaining a certain signal fidelity or suppressing fluctuations, can be quite expensive in terms of metabolic resources^3, 15^, and hence potentially an area where optimization is relevant.

Discussions of signaling performance limits are often framed in terms of information theory concepts like channel capacity^16, 17^, and complemented by direct experimental estimates^18–25^. In recent years, another tool has emerged for understanding constraints on biological signal propagation: optimal noise filter theory^15, 26–30^, drawing on the classic work of Wiener and Kolmogorov (WK) in engineered communications systems^31–33^. The theory maps the behavior of a biological network onto three basic components: a signal time series, noise corrupting the signal, and a filter mechanism to remove the noise. Once the identification is made, the payoff is substantial: one can use the WK solution for the optimal noise filter function to derive closed form analytical bounds on measures of signal fidelity (like mutual information) or noise suppression (like Fano factors). These bounds depend on the network’s reaction rate parameters, allowing us to determine a minimum energetic price associated with a certain level of performance^15^. Finally, the theory specifies the conditions under which optimality can be realized in a particular network.

To date, however, there has been one major caveat: the WK theory relies on a continuum description of the molecular populations in the network, and assumes all reaction rates are linearly dependent on the differences of these population numbers from their mean values. While this may be a good approximation in certain cases (i.e. large populations, with small fluctuations relative to the mean), it certainly raises doubts about the universal validity of the bounds derived from the theory. Biology is rife with nonlinearities, for example so-called ultrasensitive, switch-like rate functions^34^ in signaling cascades. Could these nonlinear effects allow a system to substantially outperform a WK bound derived using linear assumptions? Curiously, every earlier attempt to answer this question for specific systems^26, 28^ (summarized below) has yielded the same answer: the WK bound seemed to hold rigorously even when nonlinearities and discrete populations were taken into account.

The current work shows that this is not the full story. We have found for the first time two biological examples that can be explicitly proven to violate their WK bounds: a two-level signaling cascade and a negative feedback loop. To demonstrate this, we start by describing a general theoretical framework for signaling cascades with arbitrary numbers of intermediate species (levels), with the possibility of feedback interactions between the input and output species. We show how to calculate WK bounds based on the linearized, continuum version of this system, generalizing earlier WK results for single-level systems. In order to check the validity of the WK bound, we introduce an analytical approach for calculating exact moments of the discrete stationary probability distribution of molecular populations, starting from the underlying master equation. Our method works for arbitrarily long cascades in the absence of feedback. It allows us to find cases in a nonlinear two-level signaling cascade where the WK bound holds, as well as cases where it is violated. A similar picture emerges in a nonlinear single-level system with negative feedback, but here we use an alternative numerical approach to tackle the master equation. Remarkably, for the cases where nonlinearity helps beat the WK bound, the magnitude of the violation is tiny, typically fractions of a percent. We observe a trend that as the signaling efficiency increases, improving the biological function of the system, the size of the violation decreases or vanishes. This makes the WK value an excellent estimate for the actual performance limit in the biologically relevant parameter regime. Thus while the results show the WK theory does not rigorously bound the behavior of nonlinear signaling systems, they also put the theory on a more solid foundation for practical applications.

## 2 Results

### 2.1 Signaling network

We begin by defining a general model of an *N*-level cellular signaling cascade. Each specific system we consider in our analysis will be a special case of this model. As shown schematically in Fig. 1, we have an input chemical species *X*_0_ followed by *N* downstream species *X*_1_, …, *X*_*N*_ . For example, if this was a model of a mitogen-activated kinase (MAPK) cascade^35^, the input *X*_0_ would be an activated kinase, which activates another kinase via phosphorylation (*X*_1_), which in turn leads to a sequence of downstream activations until we reach the final activated kinase *X*_*N*_ . The copy number of species *X_i_* is denoted by *x_i_* = 0, 1, 2, Hence the state of the system can be represented by the vector ***x*** = (*x*_0_, *x*_1_, … , *x*_*N*_). Stochastic transitions between states are governed by an infinite-dimensional Markovian transition rate matrix *W*. The element *W_**x**_′,**x*** of this matrix represents the probability per unit time to observe state ***x***′ at the next infinitesimal time step, given that the current state is *x*. The values of these elements will depend on the rates of the chemical reactions that are possible in our signaling network, as described below. The probability *p*_***x***_(*t*) of being in state ***x*** at time *t* evolves according to the corresponding master equation^36^,

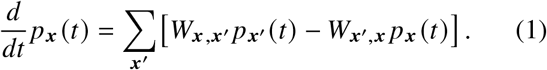

**Figure 1.**
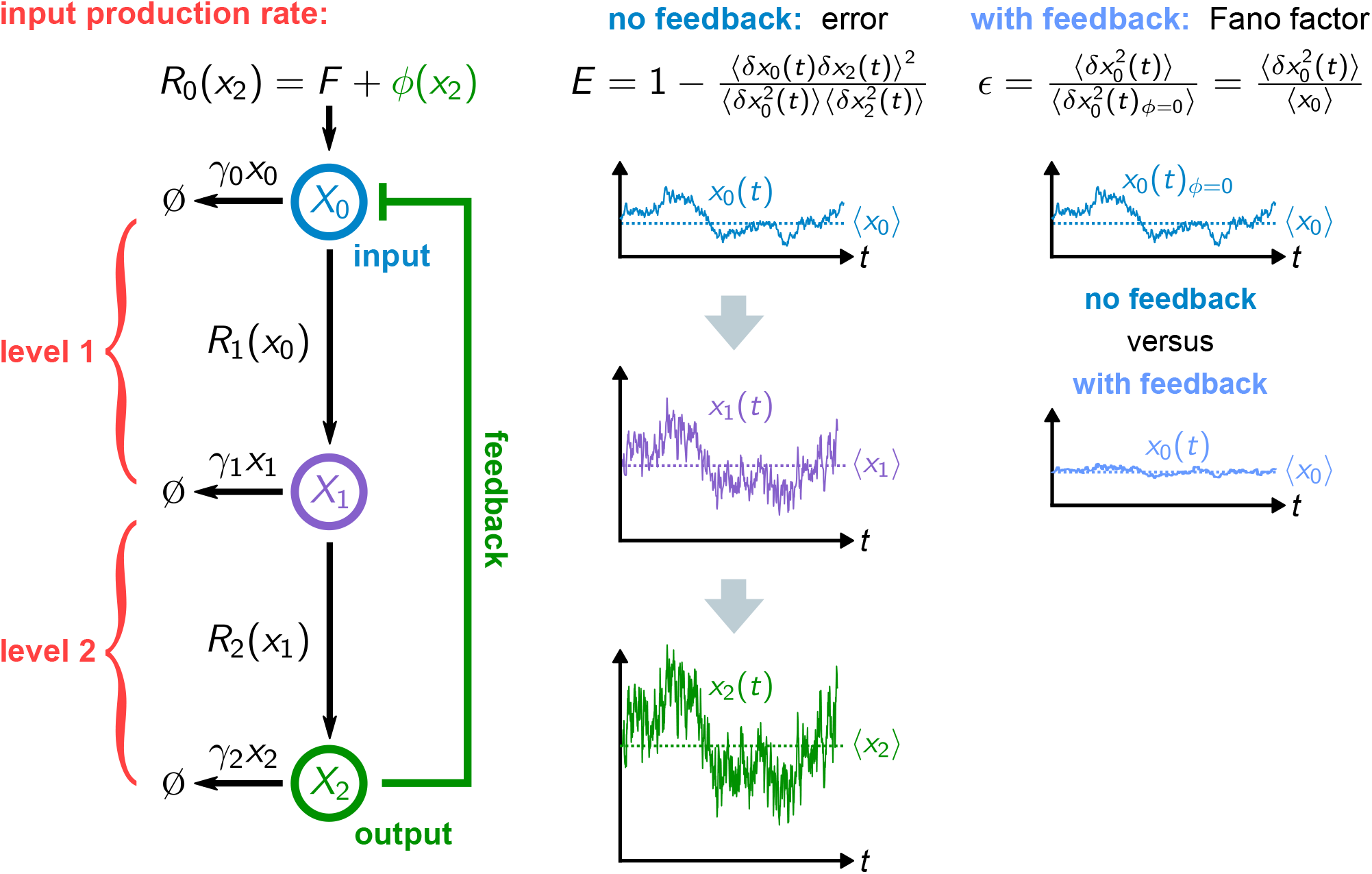
Overview of the *N*-level signaling cascade model, showing an example with *N* = 2. The signal from input species *X*_0_ is propagated through to output species *X_N_*, with the possibility of feedback back to the input. In the absence of feedback, the signal fidelity is measured via the error *E*, defined in terms of correlations between the input fluctuations *δx*_0_(*t*) = *x*_0_(*t*) – ⟨x_0_⟩ and output fluctuations *δx_N_*(*t*) = *x_N_(t) – ⟨x_N_*⟩. In the linearized system the error is related to the input-output mutual information 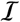 through 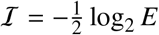. For the system with feedback, the quantity we focus on is *ϵ*, the ratio of input variance 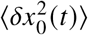 with feedback to the variance 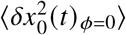 without. This is also equal to the Fano factor 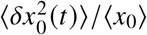, which measures the effectiveness of feedback in suppressing input fluctuations.

The first term on the right represents the gain of probability in state ***x*** due to transitions out of all other states ***x***′ into ***x***, and the second term the loss due to transitions out of ***x*** into all other states. We will focus on systems where *W* is time-independent and the system reaches a unique stationary distribution 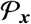. The latter satisfies Eq. (1) with the left-hand side set to zero,

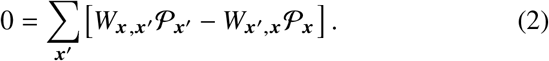

All physical observables we consider can be expressed as averages over this stationary distribution. If *f*(***x***) is some function of state ***x***, we will use 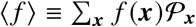 to denote the associated stationary average.

The detailed form of Eq. (2) for our cascade requires specifying all the possible chemical reactions in our network. We start with species *X*_0_, which is produced with some rate *R*_0_(*x_N_*) ≥ 0. We treat “production” as occurring with a single effective rate, encompassing all the substeps involved in activation of *X*_0_ from some inactive form (not explicitly included in the model). The functional form of the rate *R*_0_(*x_N_*) can be decomposed into two parts,

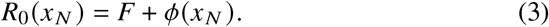

Here, *F* represents a constant baseline activation rate and *ϕ*(*x*_*N*_) the perturbation to that rate due to feedback from the final downstream species *X*_*N*_. *ϕ*(*x*_*N*_) is a potentially nonlinear function, with *ϕ*′ *x*_*N*_ < 0 corresponding to negative feedback (production of *X*_0_ inhibited by increases in *x*_*N*_), and *ϕ*′ (*x*_*N*_) > 0 corresponding to positive feedback (production of *X*_0_ enhanced by increases in *x*_*N*_). In the absence of feedback, *ϕ*(*x*_*N*_) = 0. The possibility of feedback from the last species to an upstream one has analogues in biological systems like the ERK MAPK pathway^37^. Of course there may be feedback to multiple upstream species (as is the case for ERK), but here we only consider one feedback interaction as a starting point for modeling.

In a similar spirit, the baseline rate *F* is a constant for simplicity, representing the net effect of processes leading to the activation of *X*_0_ that are not explicitly part of the model. There are also deactivation processes for *X*_0_ (i.e. the action of phosphatases) which we model by an overall deactivation rate *γ*_0_*x*_0_ proportional to the current population. We denote the constant *γ*_0_ as the per-capita deactivation rate. For the case of no feedback, the marginal stationary probability of the input *X*_0_ is a Poisson distribution 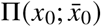^38^,

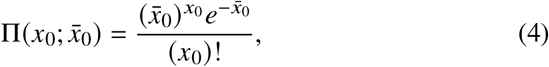

where 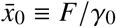, which in this case is equal to the mean and variance: 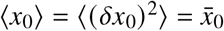, with *δx*_0_ ≡ *x*_0_ – ⟨*x*_0_⟩. Dynamically, the input signal has exponentially decaying autocorrelations, with characteristic time 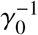. More complex types of input (for example with time-dependent *F*(*t*) or non-exponential autocorrelations) can also be considered in generalizations of the model^26, 27^. For our system, once feedback is turned on, the input distribution is no longer simply described by Eq. (4), and in general will not have a closed form analytical solution.

For *i* > 0, the production function for the *i*th species *X_i_* is *R_i_*(*x*_*i*–1_) ≥ 0, depending on the population *x*_*i*–1_ directly upstream. The deactivation rate at the *i*th level is *γ_i_x_i_*. We allow the *R_i_* functions to be arbitrary, and hence possibly nonlinear. Putting everything together, we can now write out the explicit form of Eq. (2),

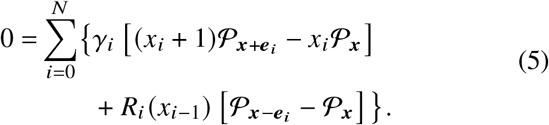

For compactness of notation, we define *x*_−1_ ≡ *x*_*N*_ and introduce the (*N* + 1)-dimensional unit vectors *e_i_*, where ***e***_0_ = (1, 0, … , 0), *e*_1_ = (0, 1, 0, … , 0), *e*_2_ = (0, 0, 1, 0, … , 0) and so on. Eq. (5) is generally analytically intractable, in the sense that we cannot usually directly solve it to find the stationary distribution 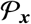. Despite this limitation, we can still make progress on understanding signaling behavior in the cascade via alternative approaches. Linearization of the production functions, described in the next section, is one such approach. Crucially, this approximation facilitates deriving bounds on signaling fidelity via the WK filter formalism. Later on we will also introduce exact analytical as well as numerical methods for tackling certain cases of Eq. (5), to explore the validity of the WK bounds in the presence of nonlinearities.

### 2.2 WK filter formalism

In this section, we provide a brief overview of linearizing our signaling model and mapping it to a WK filter, generalizing the approach developed in Refs. 26, 28, 38. This mapping allows us to derive bounds on various measures of signaling fidelity, which we know are valid at least within the linear approximation. The aim here is to summarize the bounds that we will later try to beat by introducing nonlinearities. Additional details of the WK approach can be found in the review of Ref. 38, which presents three special cases of our model: the *N* = 1 and *N* = 2 cascades without feedback, and the *N* = 1 system with feedback. The WK bound for the general *N*-level cascade, with and without feedback, is presented here for the first time, with the complete analytical derivation shown in the Supplementary Information (SI).

#### 2.2.1 Linearization

If we consider the limit where the mean copy numbers of all the chemical species in the cascade are large, we can approximately treat each population *x*_*i*_ as a continuous variable. If the magnitude of fluctuations in the stationary state is small relative to the mean, we can also approximate all the production functions to linear order around their mean values,

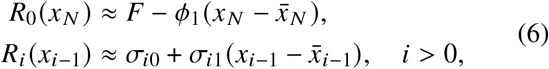

with some coefficient *ϕ*_1_, *σ*_*i*0_, and *σ*_*i*1_. Here we have absorbed the zeroth-order Taylor coefficient of *ϕ*(*x*_*N*_) around 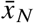 into *F*. Note that the sign convention for the first-order coefficient *ϕ*t_1_ means that *ϕ*_1_ > 0 corresponds to negative feedback. The stationary averages in the linearized case are:

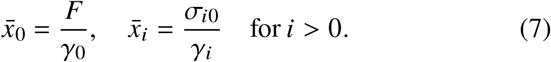

We will use bar notation like 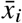 to exclusively denote the linearized stationary mean values. Brackets like ⟨*x_i_*⟩ will always denote the true mean, whether the system is linear (in which case 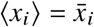) or not.

One advantage of linearization is the ability to express dynamics in an analytically tractable form, using the chemical Langevin approximation^39^. The Langevin equations corresponding to Eq. (5) are:

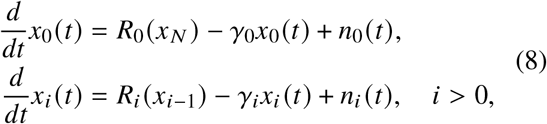

where the *n*_*i*_(*t* are Gaussian noise functions with correlations 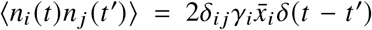 We can rewrite Eq. (8) in terms of deviations from the mean, *δx*_*i*_(*t*) ≡ *x*_*i*_(*t*) – ⟨*x*_*i*_⟩, plugging in Eqs. (6)-(7). The result is:

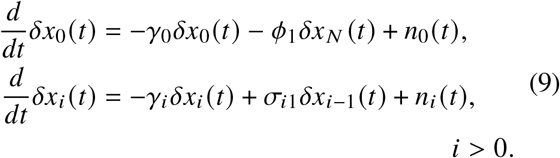

#### 2.2.2 Finding bounds on signal fidelity by mapping the system onto a noise filter

The linear chemical Langevin approach also allows us to map the system onto a classic noise filter problem from signal processing theory. We describe two versions of this mapping here, the first for the system without feedback, and the second with feedback.

##### 1. No feedback system

Imagine we are interested in understanding correlations between two dynamical quantities in our system, as a measure of how accurately signals are transduced through the cascade. The choice of these two quantities, one of which we will label the “true signal” *s*(*t*) and the other the “estimated signal” 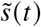 within the filter formalism, depends on the biological question we would like to ask. For the cascade without feedback (*ϕ*_1_ = 0), a natural question is how well the output *X*_*N*_ reflects the input *X*_0_. The function of the cascade can be to output an amplified version of the input^40^, but there is inevitably corruption of the signal as it is transduced from level to level due to the stochastic nature of the biochemical reactions in the network. If we assign *s*(*t*) ≡ *δ*x_0_(*t*) and 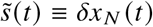, it turns out that because of the linearity of the dynamical system in Eq. (9) the two are related through a convolution,

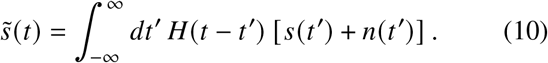

The details of the functions *H*(*t*) and *n*(*t*), as derived from Eq. (9), are given in the SI Sec. 1. One can interpret Eq. (10) as a linear noise filter: a signal *s*(*t*) corrupted with additive noise *n*(*t*) (a function which depends on the Langevin noise terms *n*_*i*_(*t*) is convolved with a filter function *H*(*t*) to yield an estimate 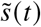. The filter function, which encodes the effects of the entire cascade, obeys an important physical constraint: *H*(*t*) = 0 for all *t* < 0. This enforces causality, since it ensures that the current value of 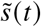 (the output in our case) only depends on the past history of the input plus noise, *s*(*t*′) + *n*(*t*′) for *t*′ < *t*.

The traditional version of filter optimization^31–33^ is searching among all possible causal filter functions *H*(*t*) for the one that minimizes the relative mean squared error between the signal and estimate:

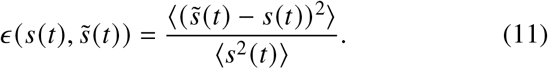

Since the averages are taken in a stationary state, *ϵ* is time-independent, and can have values in the range 0 ≤ *ϵ* < ∞. For the case of a biological cascade however, where *s*(*t*) and 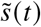 are the times series of input and output fluctuations *δx*_0_(*t*) and *δx*_*N*_(*t*) respectively, we expect that the output may be an amplified version of the input. Hence a better measure of fidelity may be a version of Eq. (11) that is independent of the scale differences between signal and estimate. To define this scale-free error, note that the optimization search over all allowable *H*(*t*) necessarily involves searching over all constant prefactors *A* that might multiply a filter function *H*(*t*). Using *AH*(*t*) as the filter function instead of *H*(*t*), is equivalent to switching from 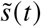 to 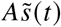, as can be seen from Eq. (10). If we were to look at the error 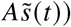 for a given 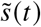 and *s*(*t*), we can readily find the value of *A* that minimizes this error, which is given by 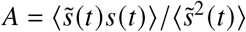. Plugging this value in, we can define a scale-free error *E* as follows:

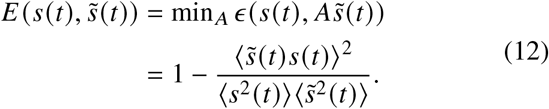

By construction, 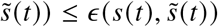 and in fact *E* has a restricted range: 0 ≤ *E* ≤ 1. The independence of *E* from the relative scale of the output versus the input makes it an attractive measure of the fidelity of information transmission through the cascade. In fact, within the linear chemical Langevin approximation, one can show that 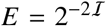, where 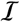 is the instantaneous mutual information in bits between *s*(*t*) and 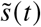^38^. Thus *E* will be the main measure of signal fidelity we focus on when we discuss the no-feedback cacade.

For a given *s*(*t*) and *n*(*t*), we denote the causal filter function *H*(*t*) that minimizes 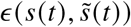 as the Wiener-Kolmogorov (WK) optimal filter *H*_WK_ *t*. Because this optimization includes exploring over all possible prefactors of *H*(*t*), the same WK filter function simultaneously minimizes 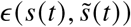 and 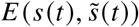, and the min-ima of the two error types coincide. We will denote this minimum as *E*_WK_. Hence we have *ϵ* ≥ *E* ≥ *E*_WK_ in general for linear systems, and *ϵ* = *E* = *E*_WK_ when *H*(*t*) = *H*_WK_(*t*).

The procedure for calculating *H*_WK_(*t*) for a specific system, and then finding the optimal error bound *E*_WK_, is based on analytical manipulation of the power spectra associated with *s*(*t*) and *n*(*t*)^33, 38^. We illustrate the details in SI Sec. 1, applying the method to our cascade model. This yields the following value for *E*_WK_ for an *N*-level cascade without feedback:

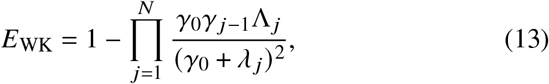

where 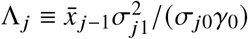 is a dimensionless parameter associated with the *j*th level, and *λ* = *λ*_*j*_ is the *j*th root with positive real part (Re(*λ*_*j*_) > 0) of the following polynomial *B*(*λ*):

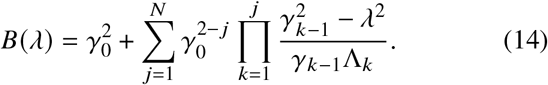

This is a polynomial of degree 2*N* in *λ*, and hence has 2*N* roots. Because the coefficients of *λ* in the polynomial are real, the conjugate of any complex root must also be a root. Finally, because only even powers of *λ* appear in *B*(*λ*), the negative of a root is also a root. Putting all these facts together ensures that there will always be *N* roots *λ*_*j*_ where Re(*λ*_*j*_) > 0. Moreover, among the set of *λ*_*j*_, any complex roots come in conjugate pairs. This guarantees that the expression for *E*_WK_ in Eq. (13) is always real. Note that the choice of ordering of the roots *λ*_*j*_, *j* = 1, … , *N* is arbitrary, since it does not affect the result. Within the linear approximation, *E*_WK_ gives a lower bound on the achievable *E*, and and hence an upper bound on the maximum mutual information between input and output, 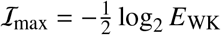.

Special cases of Eq. (13) recover earlier results. For *N* = 1, we find the single root 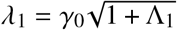, and we can rewrite Eq. (13) in a simple form:

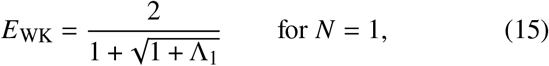

which is the result found in Ref. 26. Similarly, the *N* = 2 version, with more complicated but still analytically tractable roots *λ*_1_ and *λ*_2_, was derived in Ref. 38. For the case of general *N*, the roots *λ*_*j*_ can be found numerically. However, there is one scenario where we know closed form expressions for all the *λ*_*j*_ for any *N*. This turns out to be the case where the biological parameters of the cascade are tuned such that filter function *H*(*t*) in Eq. (10) is proportional to *H*_WK_(*t*), and hence *E* = *E*_WK_. (We do not need strict equality of the filter functions, because the resulting value of *E* is independent of an overall constant in front of *H*(*t*).) That this is even possible is itself non-trivial; generally when we vary biological parameters in a system mapped onto a noise filter, we allow *H*(*t*) to explore a certain subspace of all possible filter functions. It is not guaranteed that any *H*(*t*) in that subspace will coincide with *H*_WK_(*t*) up to a proportionality constant. However, as shown in SI Sec. 1, for the no-feedback *N*-level model we can achieve *H*(*t*) ∝ *H*_WK_(*t*) when the following conditions are met:

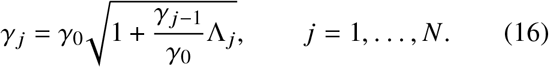

These can be solved recursively to give nested radical forms:

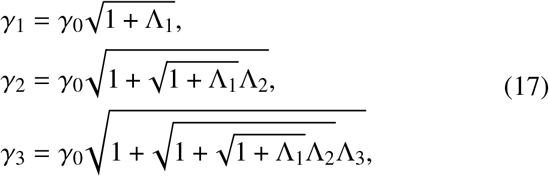

and so on. When these conditions are satisfied, the roots *λ*_*j*_ have straightforward analytical forms, namely *λ*_*j*_ = *γ*_*j*_ for all *j*. Hence we can substitute the values in Eq. (17) for *λ*_*j*_ in Eq. (13) to get *E*_WK_ explicitly when this scenario is true. With the aid of the recursion relation in Eq. (16), we can then write *E*_WK_ in this case as:

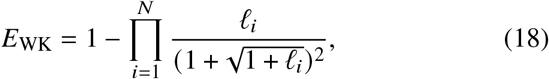

where *l*_*i*_ ≡ *γ*_*i*−1_Λ_*i*_/*γ*_0_ are dimensionless positive constants. The simple form of the bound in Eq. (18) makes it useful for analyzing the energetic cost of increasing signal fidelity in a cascade. The biological implications of this bound are discussed later on.

##### 2. System with feedback

The case with feedback uses a qualitatively different, and more abstract, mapping of the system onto a noise filter. Here the true and estimated signals are identified with the following quantities^28, 38^:

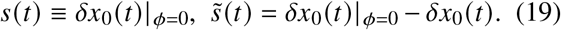

The subscript in *δx*_0_(*t*)|_*ϕ* =0_ denotes that this *δx*_0_(*t*) is obtained by solving Eq. (9) with the feedback turned off, *ϕ*(x_*N*_) = 0 or equivalently *ϕ*_1_ = 0. The *δx*_0_(*t*) without the subscript represents the solution with the feedback present. With this mapping, the error *ϵ* from Eq. (11) can be written as:

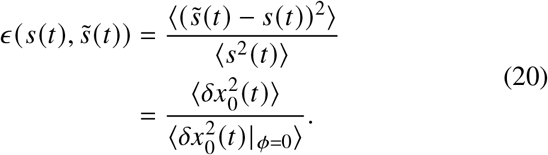

The underlying motivation is that negative feedback can serve as a homeostasis mechanism, dampening fluctuations *δx*_0_(*t*) in the *X*_0_ species that are the direct target of the feedback. Achieving a small *ϵ*, by making 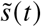 as close as possible to *s*(*t*), translates to an efficient suppression of *X*_0_ fluctuations (relative to their undamped magnitude in the absence of feedback). Note that in this case *ϵ*, rather than the scale-free error *E*, is the quantity used to specify system performance. Despite this difference, the problem is still a question of accurate information propagation through the cascade, because we need *δx*_*N*_(*t*) to encode a faithful representation of the input fluctuations in order to be able to effectively suppress them via negative feedback. Since the *X*_0_ fluctuations in the no-feedback system follow the Poisson distribution of Eq. (4), the denominator in Eq. (20) is given by 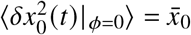. Thus 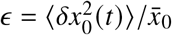, which is also known as the Fano factor (ratio of variance to the mean), a standard measure for the size of fluctuations. Poisson distributions have Fano factors *ϵ* = 1, but negative feedback in optimal cases can reduce *ϵ* to values much smaller than 1.

The close connections between the no feedback and feedback analysis is apparent when we consider the analogue of the convolution in Eq. (10) for the feedback case. It turns out that *s*(*t*) and *n*(*t*) have the same functional forms as in the no feedback case, but the filter function *H*(*t*) is different (details in SI Sec. 2). Because the WK bound depends only on the power spectra of *s*(*t*) and *n*(*t*), the result for the bound *E*_WK_ is exactly the same as Eq. (13), with roots *λ*_*j*_ specified by Eq. (14). The interpretation of Eq. (13) in this case is as a lower bound for the error in Eq. (20), namely *ϵ* ≥ *E*_WK_.

Unlike the no-feedback cascade, where we can in principle tune the biological parameters so that *E* = *E*_WK_, for the linearized negative feedback system we can only asymptotically approach the bound from above, *ϵ* → *E*_WK_. This limit is easiest to describe in the case where the production functions at each level are directly proportional to the upstream species, *R*_*i*_(*x*_*i*–1_) ∝ *x*_*i*−1_ for *i* > 0. In terms of Eq. (6), this corresponds to setting 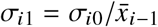, so that *R*_*i*_(*x*_*i*−1_) = *σ*_*i*1_*x*_*i*−1_. The following two conditions are then needed to approach WK optimality: i) the levels in the cascade have fast deactivation rates relative to the inverse autocorrelation time of the input, *γ*_*i*_ ≫ *γ*_0_ for *i* > 0; ii) the coefficient of the negative feedback function is tuned to the value,

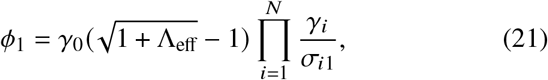

where

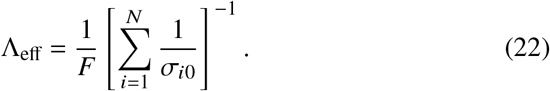

In this limit *ϵ* approaches *E*_WK_, with Eq. (13) evaluating to the same form as Eq. (15), except for Λ replaced by Λ_eff_,

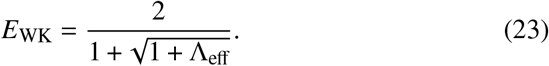

The *N* = 1 special case of this result, where Λ_eff_ = Λ_1_ = *σ*_*i*0_/*F*, was the focus of Ref. 28.

#### 2.2.3 Optimal bounds and metabolic costs

The general *E*_WK_ bound in Eq. (13), and its corresponding values in various special cases (Eqs. (15), (18), (23)), depend on the production rate parameters *σ*_*i*0_, *σ*_*i*1_ and the per-capita deactivation rates *γ*_*i*_ at each state *i*. These processes have associated metabolic costs. If production involves activation of a substrate via phosphorylation, the cell has to maintain a sufficient population of inactive substrate and also consumes ATP during phosphorylation. Similarly, deactivation requires maintaining a population of phosphatases. Achieving systems with better optimal performance can be expensive. To illustrate this, consider production functions of the form *R*_*i*_(*x*_*i*−1_ = *σ*_*i*1_*x*_*i*−1_, with 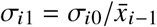, as described in the previous section. Since *ϵ* from Eq. (20) is given in terms of relative variance, a 10-fold decrease in the standard deviation of fluctuations would require a 100-fold decrease in *ε*. To decrease the optimal *E*_WK_ from Eq. (23) by a factor of 100, one would need rougly a 10^4^ increase in Λ_eff_, assuming we are in the regime where Λ_eff_ ≫ 1. This extreme cost of eliminating fluctuations via negative feedback^3^ has to be borne across the whole cascade: since Λ_eff_ in Eq. (21) is potentially bottlenecked by one *σ*_*i*0_ much smaller than the others, the mean production rates for all the levels must be hiked up in order to increase Λ_eff_.

An analogous story emerges when we analyze the same system without negative feedback. The relevant measure here is the scale-free error *E* between the time series of input and output populations, or equivalently the mutual information 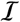. Imagine we would like to increase the mutual information upper bound 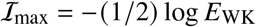 by 1 bit. In the limit of *l*_*i*_ ≫ 1 in Eq. (18), this can be achieved for example by increasing every *l*_*i*_ by a factor of ≈ 16, regardless of *N*. Given 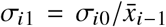 and 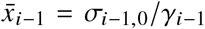 we can evaluate the dimensionless constants associated with level *i* as 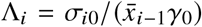 and *l*_*i*_ = (*γ*_*i*−1_/*γ*_0_)Λ_*i*_ = (*σ*_*i*0_/*σ*_*i*−1,0_)(*γ*_*i*−1_/*γ*_0_)^2^. Hence increasing *l_i_* requires either increasing the relative mean production between the *i*th level and its predecessor, or the per-capita deactivation rate of the latter (if *i* > 0), or some combination of both. Note the Λ_*i*_ parameter has a simple physical interpretation here: the average number of *i* molecules produced per molecule of *X*_*i*−1_ during the characteristic time interval 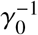 of input fluctuations. The massive cost of achieving multiple bits of mutual information between input and output in a biological signaling cascade is consistent with the narrow range of experimentally measured 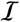 values, spanning ~ 1 to 3 bits^18–25^, with most systems near the lower end of the spectrum.

### 2.3 Nonlinearity in *N* = 1 signaling models: earlier attempts to go beyond the WK limit

The linearized noise filter approach described above provides a general recipe for deriving bounds on signaling: i) start with a linear chemical Langevin description of the system; ii) identify signal and estimate time series that are based on observables of interest, and are related via convolution in terms of some system-specific filter function *H*(*t*); iii) derive the optimal filter function *H*_WK_(*t*) and the the corresponding error bound *E*_WK_; iv) explore if and under what conditions the system can reach optimality. But the procedure leaves open an important question: is the resulting bound *E*_WK_ a useful approximation describing the system’s performance limits, or can biology potentially harness nonlinearity to enhance performance significantly beyond the WK bound? We know that nonlinear, Hill-like functional relationships are a regular feature of biological signaling^41^, manifested in some cases as an extreme switch-like input-output relation known as ultrasensitivity^34^. Is *E*_WK_ still relevant in these scenarios? This section summarizes previous efforts to answer this question (all for the *N* = 1 case), setting the stage for our main calculations.

#### 2.3.1 Nonlinearity in the *N* = 1 model without feedback

Ref. 26 derived an exact solution for the no-feedback *N* = 1 system with an arbitrary production function *R*_1_(*x*_0_) and discrete populations. The input signal remains the same as in the linear case, governed by production rate *R*_0_ (*x*_*N*_) = *F* and deactivation rate *γ*_0_. The starting point is expanding *R*_1_ (*x*_0_) in terms of a series of polynomials,

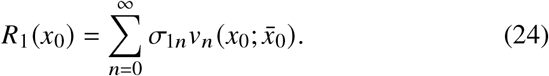

Here, 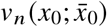 is a polynomial of *n*th degree in *x*_0_, which depends on 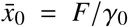 as a parameter. The functions 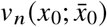 are variants of so-called Poisson-Charlier polynomials, whose properties are described in detail in the SI Sec. 4. Similar expansions have found utility in spectral solutions of master equations^42, 43^. The most important characteristic of these polynomials is that they are orthogonal with respect to averages over the Poisson distribution 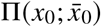 defined in Eq. (4). If we denote 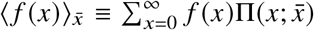 the average of a function *f*(*x*) with respect to a Poisson distribution 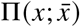, then^26, 44^

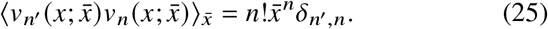

The first few polynomials are given by:

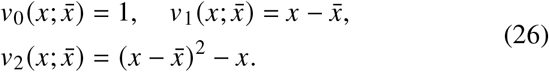

Eq. (25) allows the coefficients *σ*_1*n*_ from Eq. (24) to be evaluated in terms of moments with respect to 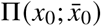,

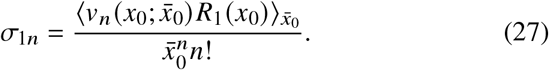

Using Eq. (26) we can write the first two coefficients as

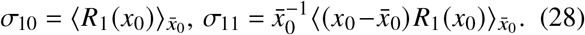

They have a simple physical interpretation: *σ*_10_ is the mean production rate and *σ*_11_ is a measure of how steep the production changes with *x* near 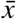, and they are exactly the same as the coefficients in the linear expansion of Eq. (6). If *σ*_1*n*_ ≠ 0 for any *n* ≥ 2 then the production function *R*_1_(*x*_0_) is nonlinear.

The exact expression for the error *E* derived in Ref. 26 takes the form:

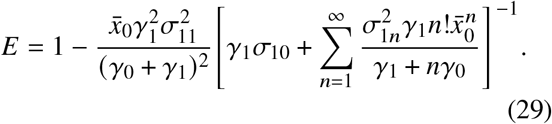

The nonlinear *σ*_1*n*_ coefficients for *n* ≥ 2 contribute to the expression in the brackets as 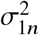 multiplying a positive factor, and hence if nonzero always act to increase the error regardless of their sign. It turns out that Eq. (29) is bounded from below by the *N* = 1 WK limit from Eq. (15),

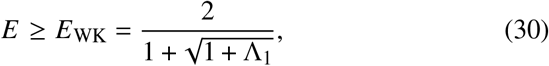

with 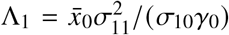. The WK limit is achieved when 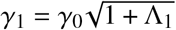, just as predicted by Eq. (17), and when *R*_1_ (*x*_0_) has the optimal linear form, 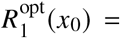 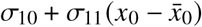, with all *σ*_1*n*_ = 0 for *n* ≥ 2.

Increasing the slope of *R*_1_(*x*_0_) at 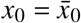 will increase *σ*_11_ and hence Λ_1_, progressively decreasing the *E*_WK_ limit. This can be seen in Fig. 2, which illustrates different production functions and the corresponding error values. Eventually, the slope will become so steep that it is impossible to have a purely linear function *R*_1_(*x*_0_) with that value of *σ*_11_. This is because *σ*_11_ must always be smaller than 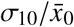 to have a linear production function that is everywhere non-negative, *R*_1_(*x*_0_) > 0 for all *x*_0_ ≥ 0. *σ*_11_ can be arbitrarily large for very steep, sigmoidal production functions *R*_1_(*x*_0_), but in this case, the error will be significantly larger than *E*_WK_ due to the contributions from the nonlinear coefficients *σ*_1*n*_, *n*≥2. We see this for the largest values of Λ_1_ in Fig. 2B, with the added error due to nonlinearity overwhelming the benefit from large Λ_1_. In summary, for the *N* = 1 no-feedback model, there is no way to beat the WK limit, regardless of the choice of *R*_1_ (*x*_0_).

**Figure 2.**
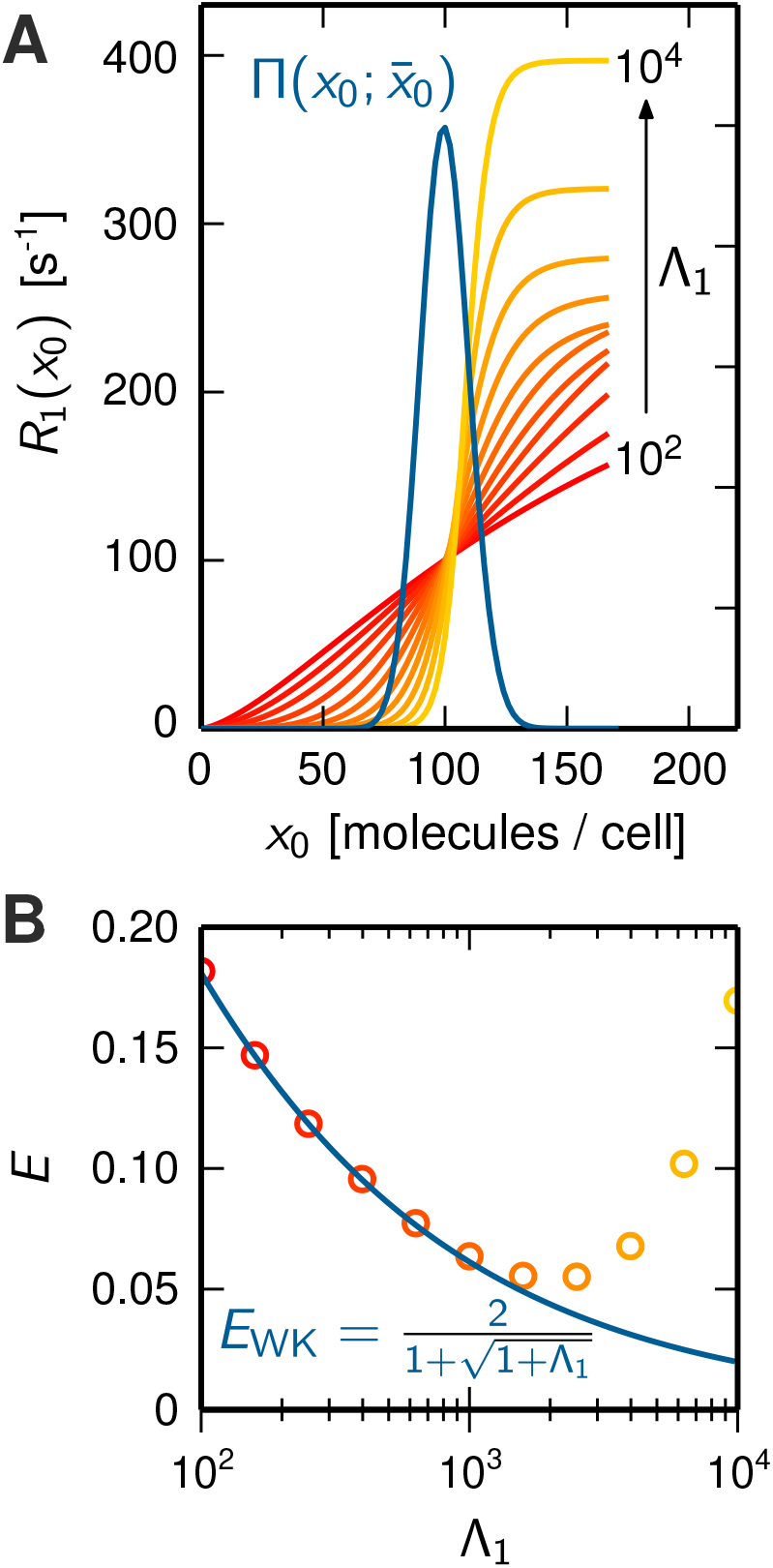
The case of a no-feedback *N* = 1 signaling model with a nonlinear production function *R*_1_(*x*_0_) for parameters: *F* = 1 s^−1^, *γ*_0_ = 0.01 s^−1^, *σ*_10_ = 100 s^−1^. A) Examples of a variety of production functions *R*_1_(*x*_0_), colored from red to yellow based on their steepness at 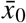, and hence the size of the corresponding parameter Λ_1_. Superimposed in black is the marginal distribution 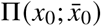 of the input species *X*_0_. B) For the production functions shown in panel A, the corresponding exact error *E* from Eq. (29) (circles) as a function of Λ_1_. The WK bound *E*_WK_ from Eq. (30) is shown in blue for comparison. Adapted from Ref. 26.

### 2.4 Nonlinearity in the *N* = 1 TetR negative feedback circuit

Ref. 28 studied an *N* = 1 negative feedback loop inspired by data from an experimental synthetic yeast gene circuit^45^. In this circuit, TetR messenger RNA (the *X*_0_ species) leads to the production of TetR protein (the *X*_1_ species), while the protein in turn binds to the promoter of the TetR gene, inhibiting the production of the messenger RNA. The model is similar to Eq. (8) when *N* = 1,

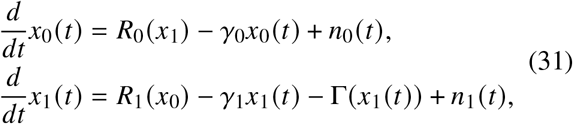

with a linear production function *R*_1_(*x*_0_) = *σ*_11_*x*_0_, but with a sigmoidal Hill function form for the feedback *R*_0_ (*x*_1_),

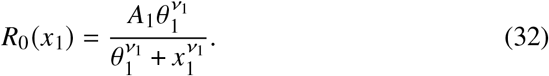

Note that this model has additional contribution to the deactivation of the output, a function Γ*x*_1_ that is also sigmoidal:

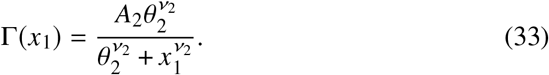

The parameters *A*_*i*_, *v*_*i*_, *θ*_*i*_, *i* = 1, 2 are all non-negative and determine the shape of the two Hill functions, which are common phenomenological expressions for regulatory interactions in biology^41^.

Using numerical methods to solve the corresponding master equation (similar to those described below), one can carry out a parameter search to solve the following optimization problem: with the constraints of fixed *γ*_0_, *γ*_1_, *σ*_11_, 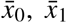 (and hence also fixed Λ_1_ = *σ*_11_/*γ*_0_), one can vary the Hill function parameters to find the smallest possible *ϵ*. The circles in Fig. 3 show the optimization results for different Λ_1_ in the range Λ_1_ = 2 – 10, comparable to experimental estimates^46^. The *E*_WK_ bound from Eq. (23) is shown for comparison as a solid curve. The dashed curve shows an exact bound for the system, derived by Lestas, Vinnicombe, and Paulsson (LVP)^3^ using information theory, which applies when *R*_1_(*x*_0_) is linear and where the negative feedback from *X*_1_ back to *X*_0_ can occur via any function (linear or nonlinear). This exact bound is given by

**Figure 3.**
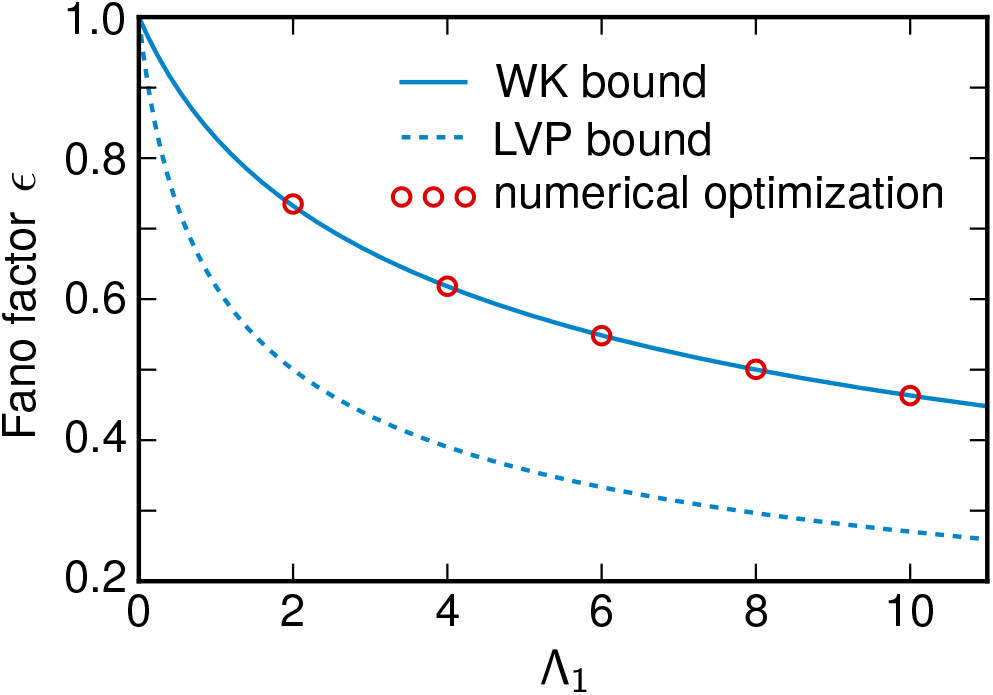
The Fano factor *E* for the TetR *N* = 1 negative feedback loop of Ref. 28. Numerical optimization results are shown as circles, while the WK and LVP lower bounds (Eqs. (23) and (34) respectively) are shown as solid and dashed curves. Figure adapted from Ref. 28.

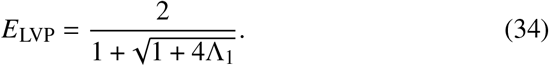

Note that *E*_LVP_ ≤ *E*_WK_, which opens the possibility that a nonlinear system could fall somewhere between the two curves, beating *E*_WK_. However, as we see in Fig. 3, the optimal numerical results were only able to approach from above, and never outperformed, the WK limit.

Based on the nonlinear *N* = 1 results with and without feedback described above, one could plausibly imagine that WK theory somehow provides universal bounds. Despite the fact that the WK limit was derived for linear systems, it surprisingly gives a rigorous bound for the nonlinear no-feedback model, and a numerical search to beat the limit proved fruitless in the feedback case. However, as we demonstrate in the next section, such a conclusion is premature.

## 3 Beating the WK limit

To explore the validity of the WK limit more broadly, we need to be able to obtain precise error results in a wider range of nonlinear signaling systems. In this section, we provide two lines of evidence that demonstrate for the first time error values below *E*_WK_. The first is from an *N* = 2 no-feedback cascade with a linear *R*_1_(*x*_0_) and quadratic *R*_2_(*x*_1_) production functions. (We will prove that the related case, where *R*_1_(*x*_0_) is nonlinear but *R*_2_(*x*_1_) linear, always gives *E* ≥ *E*_WK_.) These results use an exact expression for *E* that is valid for any *N* > 1 no-feedback system, based on a recursion relation derived from the master equation of Eq. (5) that can be evaluated numerically to arbitrary precision. The second line of evidence is from a *N* = 1 negative feedback loop with linear production function *R*_1_(*x*_0_) and a feedback function *ϕ*(*x*_1_) that includes a quadratic contribution. Here a solvable recursion relation for *E* is not possible, so we use a numerical solution of Eq. (5).

### 3.1 Exact calculation of error in the nonlinear, discrete *N* > 1 model without feedback

In order to understand the behavior of more complex nonlinear signaling cascades, we need to generalize the exact *N* = 1 no-feedback error expression from Eq. (29) to systems with *N* > 1. To start, let us introduce some convenient notation to deal with multiple-level systems. Along with the *N* + 1-dimensional vector *x* = *x*_0_, *x*_1_, … , *x*_N_ that describes the full state of our system, we will define t he *N*-dimensional truncated vector 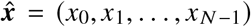 that is missing the final component *x*_*N*_. In a similar way we define the truncated *N*-dimensional unit vectors 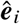, *i* = 1, … , *N* – 1, where 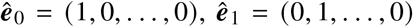, and so on until 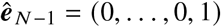. Consider the following generating function derived from the stationary distribution 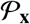,

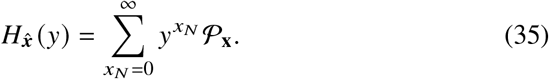

The subscript 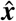 denotes the fact that 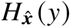 depends on all components *x*_0_ through *x*_*N*−1_, but *x*_*N*_ has been eliminated through the sum. If one carries out the sum over *x*_*N*_ on both sides of Eq. (5), one can rewrite the master equation entirely in terms of generating functions,

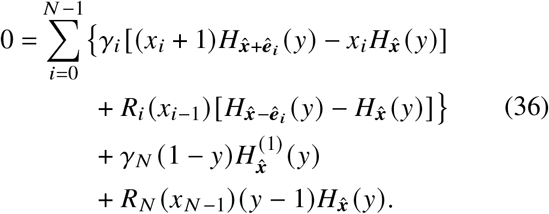

Here, the *p*th derivative of 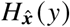 with respect to *y* is denoted as 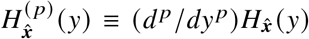. If we take *p* derivatives with respect to *y* of both sides of Eq. (36), and then set *y* = 1, we get the following relation,

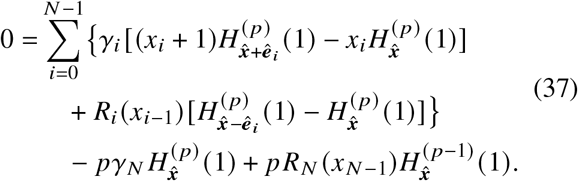

The above relation turns out to be the main one we need to evaluate the scale-free error *E*. To see this, let us rewrite the expression for *E* from Eq. (12) with the noise filter mapping 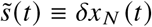, *s*(*t*) ≡ *δx*_0_ (*t*):

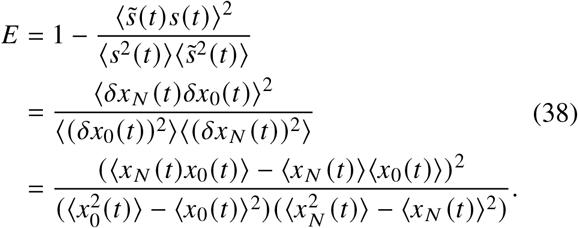

All the moments on the right-hand side of Eq. (38) with respect to the stationary distribution that involve *x*_*N*_ (*t*) can in fact be expressed in terms of 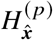 (1):

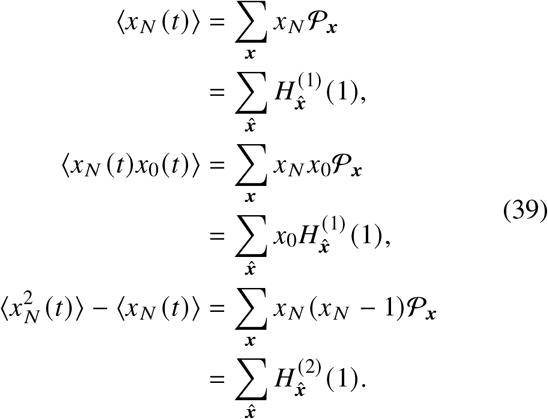

Here, 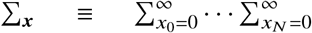 and 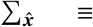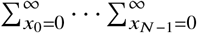. The remaining moments in Eq. (38), those that involve only *x*_0_(*t*), are known from the fact that the marginal distribution of the input *X*_0_ is just the Poisson distribution 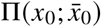 of Eq. (4). This yields

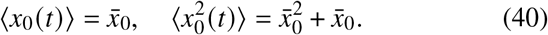

Recall that the barred notation denotes the linearized stationary averages defined in Eq. (7). Thus the approach to finding *E* is as follows: i) use Eq. (37) to derive properties of 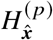 1 that allow us to evaluate the moments in Eq. (39); ii) together with Eq. (40), we can then plug the moment results into Eq. (38) to derive an expression for *E*. Here we will summarize the final result, with the full details of the derivation shown in the SI Sec. 3.

To facilitate the solution, we expand the production function in terms of Poisson-Charlier polynomials, just as in Eq. (24) for the *N* = 1 case,

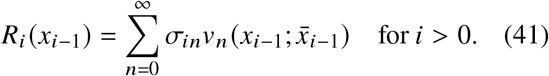

Each expansion coefficient is given by the analogue of Eq. (27), averaging over a Poisson distribution:

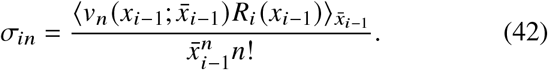

To tackle the 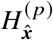 (1), we will define new functions 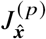 through the relation:

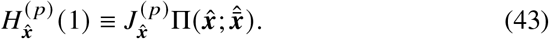

Here 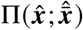 is a multi-dimensional Poisson distribution,

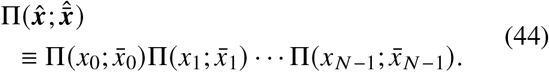

Thus any 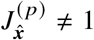 represents the deviation of 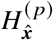 (1) from a simple multi-dimensional Poisson distribution. Similarly, we can define a multi-dimensional version of the Poisson-Charlier polynomials,

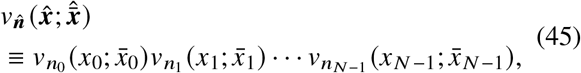

where 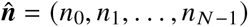 is an *N*-dimensional vector of integers *n*_*i*_ ≥ 0. Let us expand 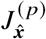 these polynomials,

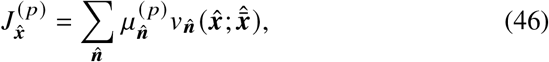

defining expansion coefficients 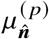. It turns out the moments in Eq. (39) are all just linear combinations of the 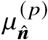, which follows from the properties of the Poisson-Charlier polynomials averaged with respect to Poisson distributions:

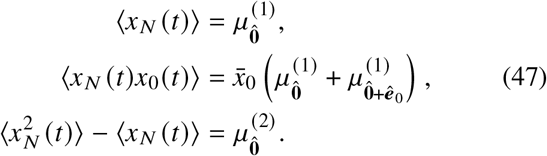

Here 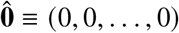is the *N*-dimensional zero vector. Plugging this into Eq. (38) gives

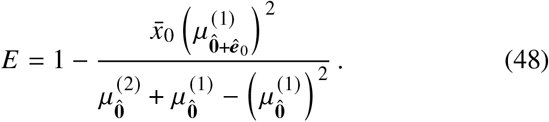

The final piece of the solution is converting Eq. (37) into a recursion relation for the coefficients 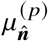:

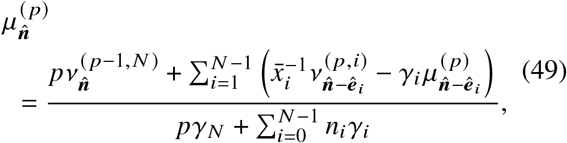

with 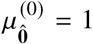. The coefficients ^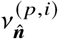^ are given by the following expansion in terms of *σ*_*in*_ and 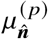:

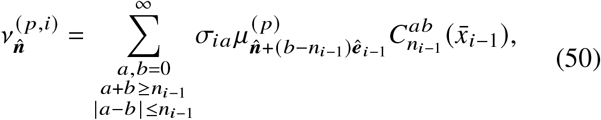

and 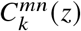 are polynomials in *z* given by:

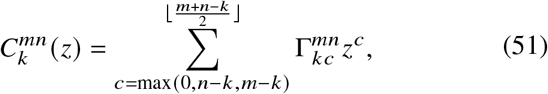

with

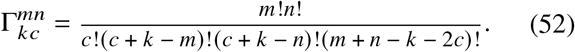

Here the sum starts at the largest of the three values 0, *n* – *k*, or *m* – *k*, and ⌊*w*⌋ denotes the largest integer less than or equal to *w*.

The general procedure for calculating *E* works as follows:

1. For a given set of production functions *R*_*i*_(*x*_*i*–1_, we calculate the expansion coefficients *σ*_*in*_ using Eq. (42). If necessary, we truncate the expansion above some order *M*, setting *σ_*in*_* = 0 for *n* > *M*. In practice, because of the rapid convergence of Eq. (41) for *x*_*i*−1_ near 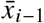, choosing *M* = 3 or 4 is sufficient. But we can increase the cutoff *M* to get whatever numerical precision we desire.
2. We plug the resulting *σ*_*in*_ into Eq. (50), and this in turn defines the 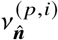 that appear in Eq. (49).
3. We solve the recursive system of equations in Eq. (49) for 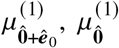, and 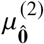, and use these to find *E* from Eq. (48).

Though complex in appearance, the procedure is easy to implement as a numerical algorithm, and with any finite cutoff *M* is guaranteed to yield a value for *E*. As we increase *M* we generally quickly converge to the exact *E* for the system.

In some cases, the entire procedure can be carried out analytically to give exact closed form expressions for *E*. When *N* = 1, we recover the result in Eq. (29), as expected. Another example is the *N* = 2 system where the first level production function *R*_1_ (*x*_0_) is arbitrary, but the second level function *R*_2_ (*x*_1_) is linear (and hence *σ*_2*n*_ = 0 for *n* ≥ 2). Here *E* is given by:

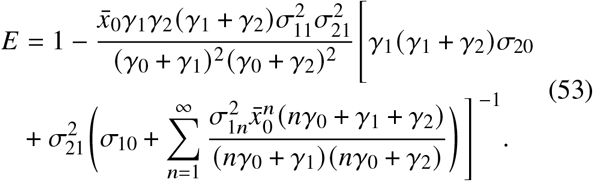

Just as in Eq. (29), any nonlinear contributions to *R*_1_ (*x*_0_) always increase *E*, since the coefficients *σ*_1*n*_ for *n* ≥ 2 only appear in the brackets in Eq. (53) as 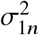 multiplying positive factors. In this scenario, *E ≥ E*_WK_ always, where *E* ≥ _WK_ is given by Eq. (13).

The simplest case where we are able to observe a violation of the *E*_WK_ limit is for *N* = 2 when *R*_1_(*x*_0_) is linear and the *R*_2_(*x*_1_) is quadratic: *σ*_1*n*_ = 0 for *n* ≥ 2 and *σ*_2*m*_ = 0 for *m* ≥ 3. The resulting analytical expression for *E* is complicated, but we can investigate its optimal behavior numerically. In Fig. 4 we conducted a numerical minimization of *E* with respect to *γ*_2_ and the quadratic coefficient *σ*_22_ for various combinations of *r* ≡ *γ*_1_/*γ*_0_ and Λ_1_, keeping Λ_2_ fixed. If we denote this minimum value *E*_min_, Fig. 4 shows log_10_|*E*_min_/*E*_WK_ – 1| with the cool colored contours indicating *E*_min_ > *E*_WK_ and warm colored contours indicating *E*_min_ < *E*_WK_. In the purely linear case described earlier, we found that *E* = *E*_WK_ when the conditions from Eq. (17) are satisfied, which corresponds to 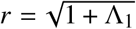 shown as as dashed white curve in the figure. With the addition of the quadratic term in *R*_2_(*x*_1_), the region near that curve now supports solutions that beat the WK limit (the warm colored band in Fig. 4). However the improvement relative to the WK bound is exceedingly small, roughly ~ 0.001 – 0.01% better.

**Figure 4.**
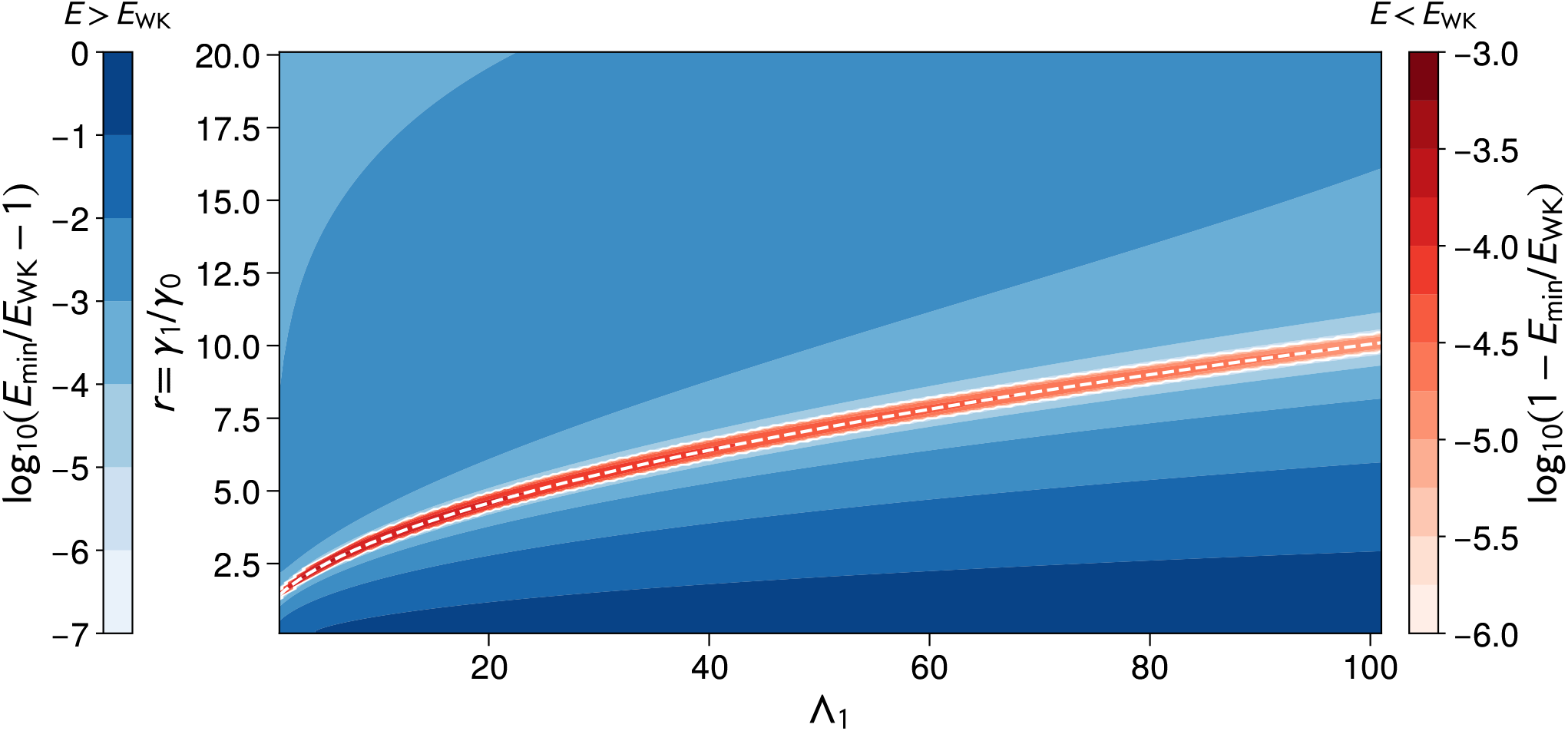
Contour plot of log_10_|*E*_min_/*E*_WK_ – 1 for the *N* = 2 cascade with linear *R*_1_(*x*_0_) and quadratic *R*_2_(*x*_1_). The minimum value of the error *E*_min_ at a given *r* = *γ*_1_/*y*_0_ and Λ_1_ is found by numerical minimization with respect to *γ*_2_ and *σ*_22_, with fixed Λ_2_ = 5. Cool colors denote regions where *E*_min_ > *E*_WK_ and warm colors where *E*_min_ < *E*_WK_. The dashed white curve corresponds to 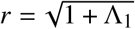.

To understand the small size of the improvement, let us look at a subset of the parameter space that is analytically tractable. Set the linear portions of the production functions to be directly proportional to the upstream population, *R*_1_(*x*_0_) = *σ*_11_*x*_0_ and 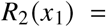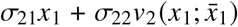, which means 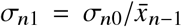 for *n* = 1, 2. Furthermore, imagine that the conditions of Eq. (17) are fulfilled f or *γ*_1_ and *γ*_2_, w hich means *E* = *E*_WK_ when the quadratic perturbation *σ*_22_ = 0. In this case Λ_1_ = *r*^2^ – 1, with *r* ≡ *γ*_1_/*γ*_0_ > 1, and Λ_2_ = *ρr*, with *ρ* ≡ *σ*_20_/*σ*_10_. *E*_WK_ from Eq. (18) can then be written as:

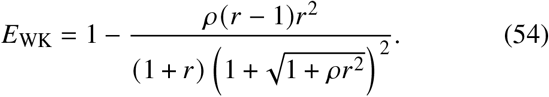

Let us focus on the regime where signaling is at least as effective as in many experimentally measured cascades^18–25^, which means *I*_max_ ≳ 1 bit or equivalently *E*_WK_ ≲ 1/4. This generally requires *r* ≫ 1 and *ρ* ≫ 1. In this limit, the complicated full expression for *E* simplifies, and we can expand the difference *E* – E_WK_ to second order in the perturbation parameter *σ*_22_,

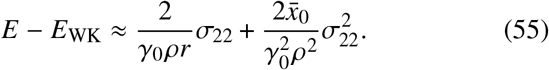

There is a minimum *E* = *E*_min_ at 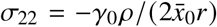, with

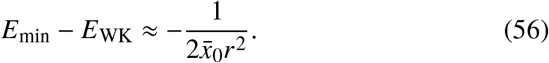

Though we can violate the *E*_WK_ bound, the size of the violation becomes small for *r* ≫ 1. From Eq. (54) we know that *E*_WK_ ~ 2/*r* for *ρ*, *r* ≫ 1, and hence the relative magnitude 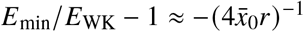. The negligible scale of the improvement over *E*_WK_ is consistent with the numerical results of Fig. 4, though the latter was calculated over a broader portion of the parameter space.

### 3.2 Revisiting nonlinearity in the *N* = 1 model with feedback

The violation of the *E*_WK_ bound in the no-feedback case raises the question of whether similar results are possible in the presence of feedback. We return to *N* = 1 system used for the TetR model above, but with several simplifications: i) we do not include the additional nonlinear degradation term Γ(*x*_1_); ii) rather than a Hill function for *R*_0_ (*x*_1_), we use Eq. (3), with a quadratic form for the feedback function *ϕ*(*x*_1_),

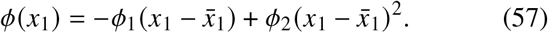

Depending on the values of *ϕ*_1_ and *ϕ*_2_, there could be a range of *x*_1_ where *R*_0_(*x*_1_) in Eq. (3) becomes negative, which is unphysical. In our numerical calculations, we thus always use max(*R*_0_ (*x*_1_), 0) as the feedback function.

However for the parameters we explored, the range of *x*_1_ where the sign switch in *R*_0_(*x*_1_) occurs is far outside the typical range of stationary state *x*_1_ fluctuations, so the precise details of the cutoff have a negligible influence on the results. A final important difference from the TetR model is that we will also investigate the regime of smaller Λ_1_ (the numerics in the earlier study were confined to Λ_1_ ≥ 2). Based on the intuition from the no-feedback case, we guess that any violation of the *E*_WK_ bound might become very small for large Λ_1_, and hence difficult to detect numerically.

Though Eq. (57) has a simple form that is convenient for parameter exploration, it has one feature that makes it somewhat unrealistic from a biological perspective. For *ϕ*_1_ > 0, *ϕ*_2_ < 0 (the case that will be of interest to us below) the slope *dϕ*(*x*_1_)/*dx*_1_ becomes positive for 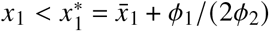, corresponding to positive feedback for smaller *x*_1_ populations. Since we would like to concentrate on systems with negative feedback, we also define an alternative feedback function 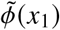 that avoids this issue by being constant for 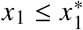 and monotonically decreasing for 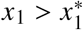:

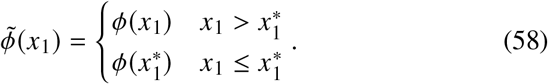

As we will see below, it turns out that both *ϕ*(y) and 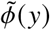 give qualitatively similar results.

The Poisson-Charlier expansion approach of the previous example can also be applied to a general *N* level feedback system, yielding a set of coupled linear equations for the coefficients 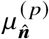 analogous to Eq. (49). However, because of the feedback interaction between *x*_*N*_ and *x*_0_, these equations are no longer particularly useful: lower order coefficients depend on higher order ones in an infinite hierarchy of equations that has no closure for any nonlinear *ϕ*(*x*_1_). We thus turn to an alternative approach: solving the master equation, Eq. (5), for the 2D stationary probability 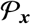, where *x* = (*x*_0_, *x*_1_). Since *x*_0_ and *x*_1_ can be any non-negative integer, Eq. (5) is an infinite linear system of equations. To make it amenable to a fast numerical solution, we truncate the range of allowable (*x*_0_, *x*_1_) to be within six standard deviations of 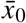 and 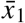. We estimate the standard deviations from the linear case (*ϕ*_2_ = 0), where closed form expressions are available in terms of the system parameters. The actual standard deviations in the presence of nonzero *ϕ*_2_ for the parameter range we considered were not perturbed significantly, so this estimation procedure worked well. Similarly, 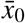 and 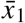 were good estimates for the actual ⟨*x*_0_⟩ and ⟨*x*_1_⟩, because the mean of the distribution shifts only a small amount with *ϕ*_2_. The window established by this procedure had a typical width of around ~ 100 for *x*_1_ and ~ 700 for *x*_0_ for parameters in the range described below. In Eq. (5), all 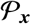 outside the allowable range of *x* were set to zero. This means that Eq. (5) becomes a finite system of linear equations that can be solved efficiently using sparse matrix methods. Once the stationary distribution is known numerically, one can then easily calculate the error *ϵ* from Eq. (20) by finding the marginal distribution of *x*_0_ and calculating its first and second moments. We checked for convergence and boundary effects by redoing the solution using window widths that were different than six standard deviations, and verified that the results were unchanged up to the desired precision (< 10^−4^ for the calculation of *ϵ*). For select parameter sets, we also validated the moments of the stationary distribution against kinetic Monte Carlo simulations^47^, though for the latter achieving high precision is difficult because of the computational time required.

We used the following parameter values (all in units of s^−1^): *γ*_0_ = 2, *γ*_1_ = 200, *σ*_10_ = 8000, *σ*_11_ = 2. The value of *F* was varied to allow for a range of possible 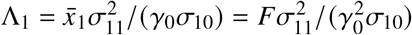. The value of *ϕ*_1_ was set to the optimality condition from Eq. (21),

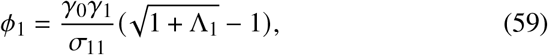

where we have used the fact that Λ_eff_ = Λ_1_ for *N* = 1. This guarantees that in the linear feedback case of *ϕ*_2_ = 0, the system should be close to the WK limit (up to correction factors due to finite *γ*_1_, since technically the WK limit is only approached in the feedback case when *γ*_1_ → ∞). Fig. 5A shows numerical results the Fano factor *ε* as a function of *ϕ*_2_ for different values of Λ_1_ between 0.25 and 1. In all cases for linear feedback (*ϕ*_2_ = 0), we see that *ϵ* > *E*_WK_, where *E*_WK_ is given by Eq. (23). The fact that *ϵ* is above *E*_WK_ for the linear system is due to the fact that *γ*_1_ is finite. For the case *ϕ*_2_ > 0 (not shown in the graphs), the error increases, while for *ϕ*_2_ < 0 we see that the error decreases, until it dips below the *E*_WK_ line before increasing again. The choice of feedback function, *ϕ*(*x*_1_) or 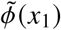, does not make a significant difference. Interestingly, the violation of the WK bound is quite small, just as in the no-feedback case, as we can see more clearly in Fig. 5B, where the ratio *ϵ*/*E*_WK_ is plotted, in this case using the 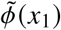 function. The largest dip we observed is still only about 1.5% below *E*_WK_. Moreover, in order to see any violation at all we had to look at small Λ_1_ ≤ 1. In this regime, *E*_WK_ is quite large, just below the Poissonian Fano factor value of 1. Hence the fluctuations are only slightly reduced by the feedback. Once Λ_1_ becomes larger, in the more biologically relevant regime where negative feedback is effective at suppressing fluctuations, we found it impossible to observe any violations of *E*_WK_. This could possibly explain the lack of any evidence of violations in the earlier TetR study^28^ (see Fig. 3), where only Λ_1_ ≥ 2 was considered. Though the figures show results for only one set of parameter values, other sets we tried produced qualitatively similar results: the nonlinear case beat the WK limit for small Λ_1_, but it was always by a small amount.

**Figure 5.**
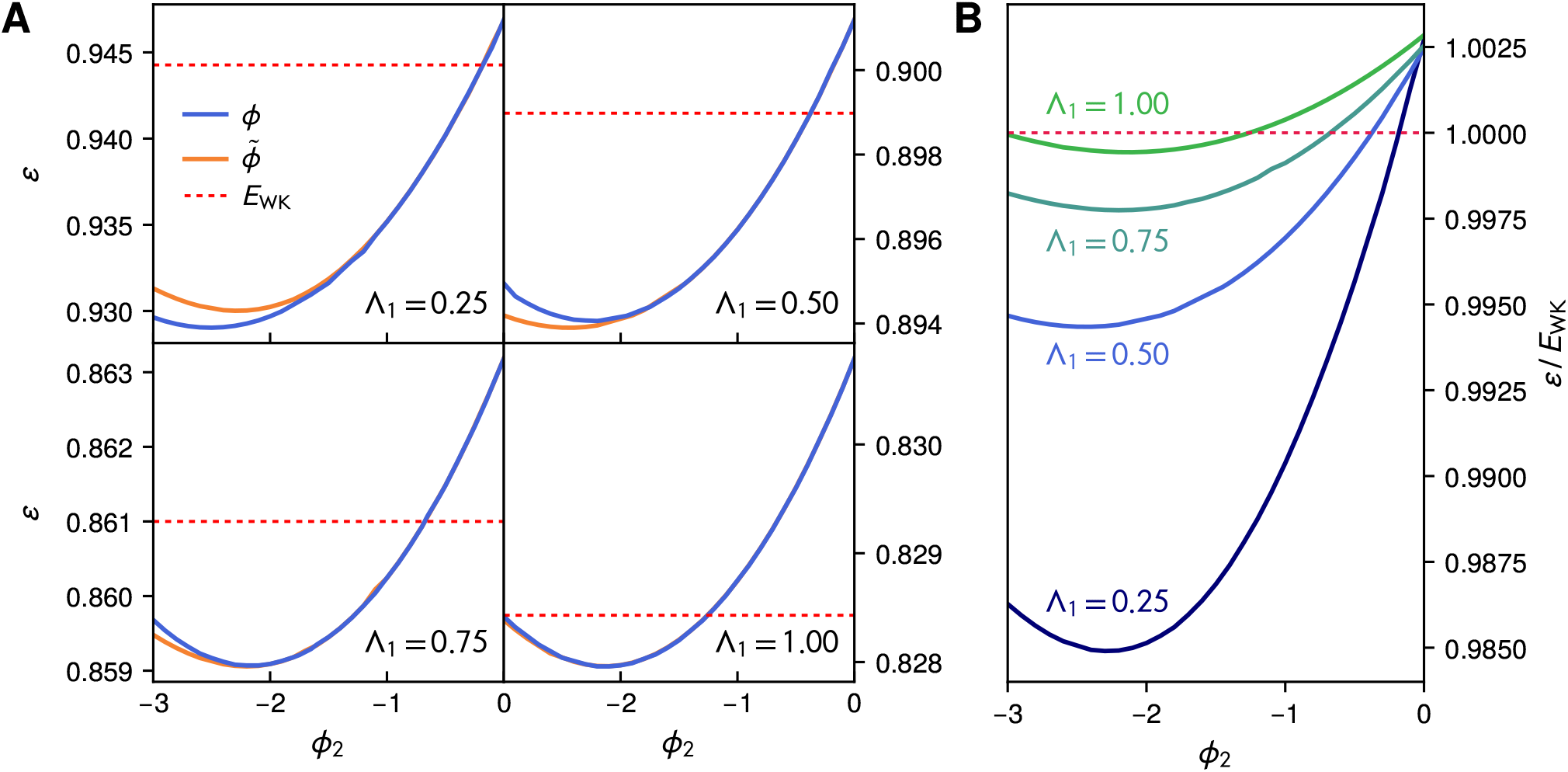
A) Numerically calculated Fano factor *ϵ* in the *N* = 1 nonlinear feedback system. The plots show *E* versus *ϕ*_2_ for the quadratic feedback function *ϕ*(*x*_1_) (blue) from Eq. (57) and the monotonic alternative 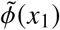 (orange) from Eq. (58), using the parameters described in the text. The WK bound *E*_WK_ in shown as a dashed red line. The subgraphs depict cases with four different values of A_1_ between 0.25 and 1. B) The Fano factor results from panel A, using the feedback function 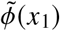, but normalized with respect to *E*_WK_. The dashed red line is *ϵ* /*E*_WK_ = 1.

## 4 Conclusions

Using a combination of analytical and numerical approaches, we have been able to show that the Wiener-Kolmogorov optimal error *E*_WK_ is not a universal lower bound for biological signaling cascades, both with and without feedback. However, far from undermining the usefulness of the WK theory, our results actually strengthen its practical value as a general purpose approximation to estimate performance limits in signaling systems. In some cases, for example the *N* = 1 or *N* = 2 no-feedback systems with nonlinear production in the first level, the *E*_WK_ bound continues to hold rigorously despite nonlinearity. And in all cases where the bound is broken, the extent of the violation is negligible and decreases or vanishes in the regime where the system is effective at its respective task (either propagating the upstream signal with high fidelity or suppressing fluctuations). Further study is needed to see if the performance gain beyond the *E*_WK_ bound can be made substantial, for example by combining the effects of nonlinearity from multiple levels in the cascade. However, additional nonlinearity is not necessarily beneficial: in Eqs. (29) and (53), and as depicted in Fig. 2, each higher order nonlinear contribution pushes us further away from the *E*_WK_ limit.

Thus for practical purposes, the WK approach remains an excellent way of deriving biological bounds that remain meaningful even when the underlying assumptions of the theory (like linearity) no longer strictly hold. Equally importantly, the theory allows one to ascertain under what conditions one can actually achieve this kind of optimality. In all the signaling systems investigated so far, *E*_WK_ is either directly attainable or can be asymptotically approached by tuning parameters. This is in contrast to a rigorous bound like *E*_LVP_ from Eq. (34), which holds for arbitrarily complex feedback mechanisms in a system with linear production. However, it has overestimated the optimal capabilities of all the feedback networks we have investigated: none of our systems ever gets close to *E*_LVP_. A recent example of the versatility of the WK theory is the study of kinase-phosphatase signaling networks in Ref. 15. A simple analytical WK bound, derived from a linearized *N* = 1 network, explains a previously unknown optimal relationship between signal fidelity, bandwidth, and minimum ATP consumption. It holds across a vast biological parameter space deduced from bioinformatic databases, and remains valid even when all the microscopic, nonlinear reaction details of the system are taken into account. The robustness of the WK bound, highlighted in the results of the current study, help us understand the theory’s success in such contexts.

Beyond future applications of WK theory to other specific systems, and possible experimental validation, there is still work to be done in developing the analytical techniques (like the Poisson-Charlier expansion) which we used for the no-feedback cascade. Exact results in nonlinear systems are relatively rare and hence valuable in themselves, and also as benchmarks for a variety of simpler approximations like the WK theory. The expansion method we described is currently limited by cases where the recursive system of equations does not close (i.e. in the presence of feedback, and more generally in biochemical networks with loops). Carefully tailored moment closure approaches^48^ might provide a way forward, and broaden the applicability of the method to systems with different types of feedback and other more complex network motifs.

## Acknowledgements

This article was submitted as part of the Dave Thirumalai Festschrift. M.H. would like to acknowledge Dave’s absolutely formative role in his own scientific career: first as a mentor and role model during my postdoctoral studies, and continuing to this day as a collaborator and friend. Discussions with Dave shaped my view of what it means to be a biophysical theorist (and so much else) and I happily see his influence live on in my own scientific mentoring of students. The work described in the current article is a coda to a line of research that first began under Dave’s auspices in 2013, when we developed the Wiener-Kolmogorov approach for biological signaling systems. The open question raised by our first article on the topic—can the WK bound ever be beaten via nonlinearity—is here answered in the affirmative. But instead of weakening the applicability of WK theory, the surprising insignificance of nonlinear enhancements actually strengthens the case for it.

## Supplementary Information

### 1 Deriving the WK optimal filter results for the multi-level cascade without feedback

#### 1.1 Mapping the system onto a noise filter

The starting point for the derivation is the system of equations in main text Eq. (9), with *ϕ*_1_ = 0 in the absence of feedback:

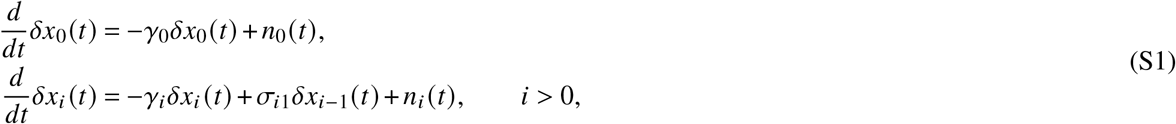

where the Gaussian noise functions satisfy 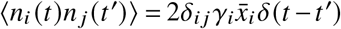). Taking the Fourier transform of Eq. (S1), we can solve the system of equations for the fluctuation functions *δx*_*i*_ (*ω*) in Fourier space,

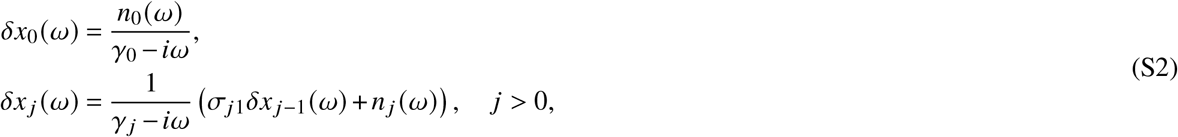

with *f*(*ω*) denoting the Fourier transform of a function *f*(*t*). Iteratively plugging the result for *δx*_*j*-1_(*ω*) into the *δx*_*j*_(*ω*) equation, starting from *j* = 1, we can solve Eq. (S2) to get the following expressions for the Fourier space input and output fluctuations:

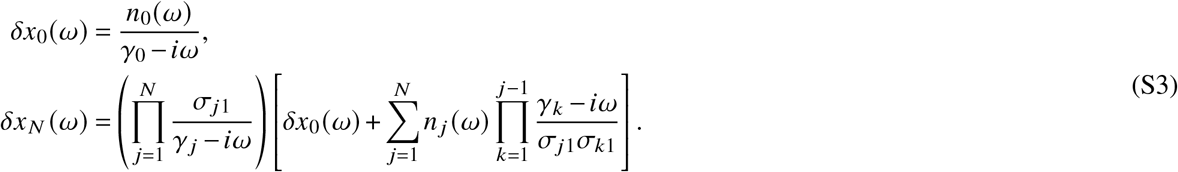

Let us compare the result for *δx*_*N*_(*ω*) to the Fourier transform of main text Eq. (10), the noise filter convolution integral:

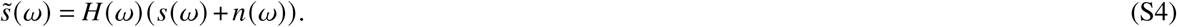

We can make a mapping of the system to a linear noise filter with the following choice of estimate, signal, noise, and filter function:

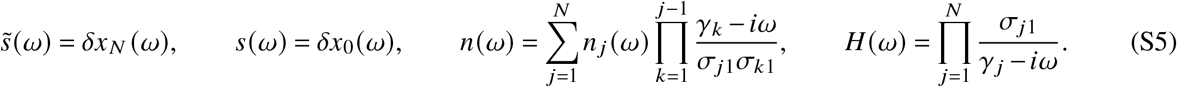

#### 1.2 Concise overview of WK optimal filter theory

To apply WK theory to our problem, let us summarize its main results (see Ref. 1 for a more detailed review). Given a Fourier-transformed signal and noise functions *s*(*ω*) and *n*(*ω*), let us denote the corresponding power spectra *P*_*s*_(*ω*) and *P*_*n*_(*ω*). The spectra are defined through the relation ⟨*f*(*ω*)*f*(*ω*′)⟩ = 2*π*P_*f*_(*ω*)*δ*(*ω* + *ω*′), where *f* = *s* or *n*. For the signal corrupted by noise, *y*(*ω*) ≡ *s*(*ω*) + *n*(*ω*), the corresponding power spectrum is *P*_*y*_(*ω*) = *P*_*s*_(*ω*) + *P*_*n*_(*ω*) if the noise is uncorrelated with the signal. This is indeed the case, since the Gaussian noise functions *n*_*j*_(*ω*) in Eq. (S5) that contribute to *n*(*ω*) are uncorrelated with *n*_0_(*ω*), the function that enters into the signal *δx*_0_(*ω*) in Eq. (S3).

Once *P*_*s*_(*ω*) and *P*_*s*_(*ω*) are specified, one can find a corresponding optimal filter function *H*_WK_(*ω*). Optimality here means that the time-domain function *H*_WK_*t*, plugged into the convolution integral of main text Eq. (10), minimizes the error 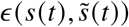 between the estimate and signal defined in main text Eq. (11). In Fourier space the optimal filter takes the following form if signal and noise are uncorrelated^2^:

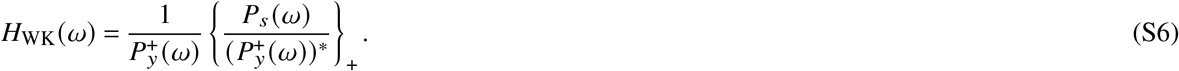

The + superscripts and subscripts denote two types of causal decompositions. For example, the function 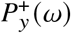 is defined via 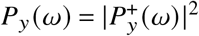, where the factor 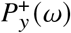 is chosen such that it has no zeros or poles in the upper half-plane. This decomposition always exists for all the physical power spectra we encounter in signaling contexts. The other decomposition, denoted by {*G*(*ω*)}_+_ for a function *G*(*ω*), can be calculated from 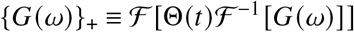. Here 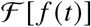 indicates the Fourier transform of a function *f*(*t*), 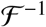 the inverse Fourier transform, and Θ(*t*) is a unit step function^3^. In practice, it is often convenient to calculate it through an alternative method: doing a partial fraction expansion of *G*(*ω*) and keeping only those terms with no poles in the upper half-plane.

To find the lower bound on *ϵ*, we inverse Fourier transform *H*_WK_(*ω*) back to the time domain. The minimum error *E*_WK_ can then be expressed compactly in the following form, which is convenient for calculations:

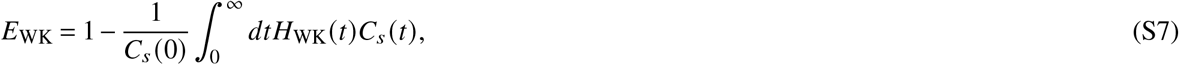

where 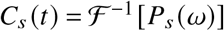 is the signal autocorrelation function, given by the inverse Fourier transform of its power spectrum.

#### 1.3 Calculating the optimal filter function *H*_WK_

Given Eqs. (S3), (S6), and the properties of the Gaussian noise functions *n*_*j*_(*t*), which in Fourier space satisfy 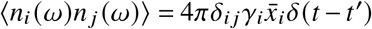, the power spectra for the signal and noise can be written as:

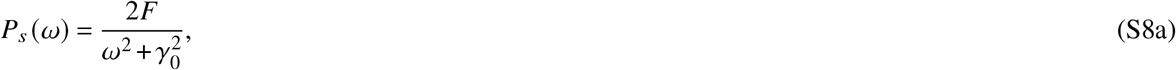

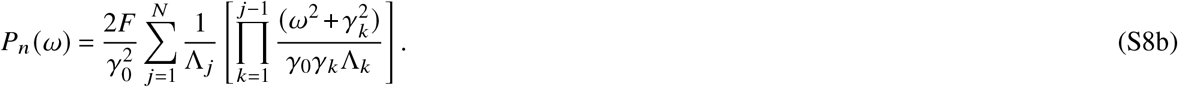

Here we have used the facts that 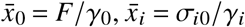 for *i* > 0, and have introduced the dimensionless constants 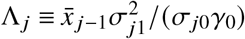. Summing *P*_*s*_(*ω*) and *P*_*n*_(*ω*), we can write *P*_*y*_(*ω*) in the form:

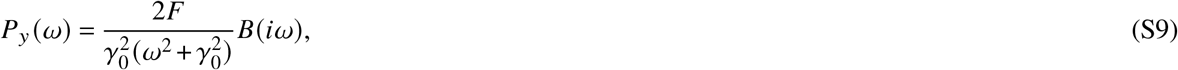

where *B*(*λ*) is the polynomial from main text Eq. (14),

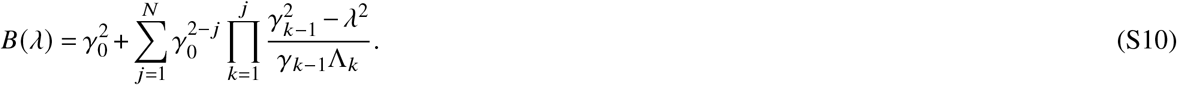

As discussed in the main text, this polynomial will always have *N* roots *λ*_*j*_, *j* = 1, … , *N*, where Re(*λ*_*j*_) > 0. (The other *N* roots of the polynomial are just −*λ*_*j*_.) Thus we can factor *B*(*iω*) in the following way:

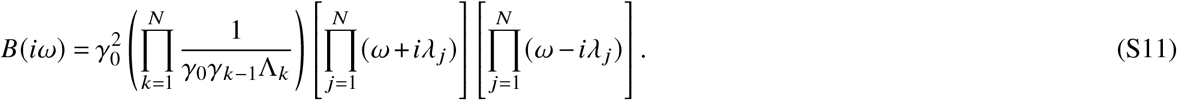

Since *ω* = *i,λ*_*j*_ for *j* = 1, … , *N* are all the zeros of *B*(*iω*) in the complex lower half plane, this enables us to write down the decomposition 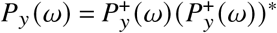 where

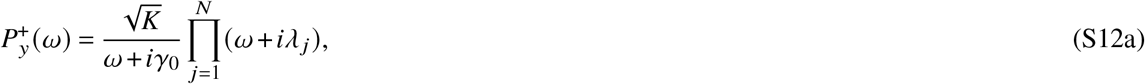

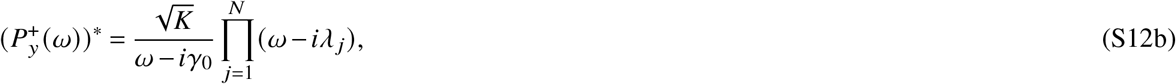

and

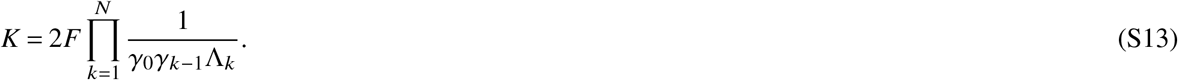

Continuing with the calculation of *H*_WK_(*ω*), we see that:

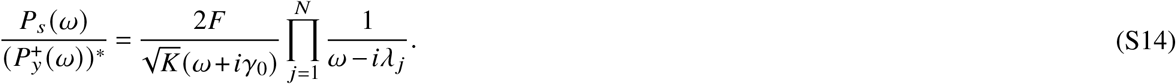

The quantity 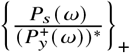 is computed from taking the causal part of the partial fraction decomposition of Eq. (S14). Because the only causal pole (pole in the lower half plane) of Eq. (S14) is −*iγ*_0_, all other terms in the decomposition are dropped, yielding:

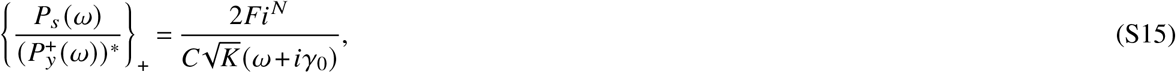

where 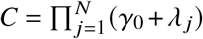. Finally, we can divide this result by 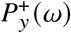, following Eq. (S6), giving us the optimal filter:

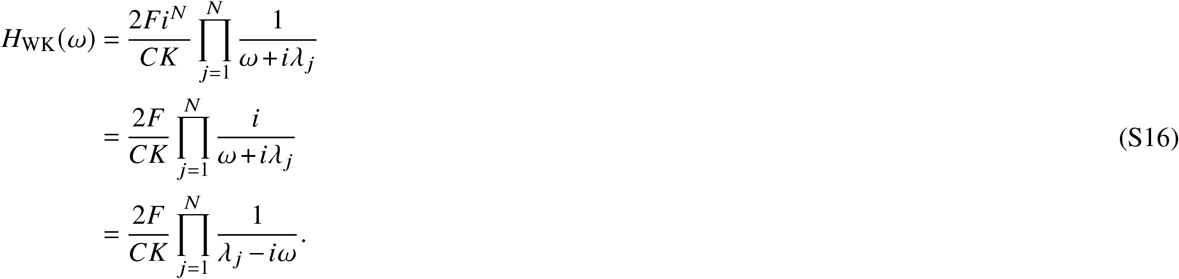

Plugging in the definitions of *C* and *K*, we can rewrite the prefactor to get the final form for the optimal filter function:

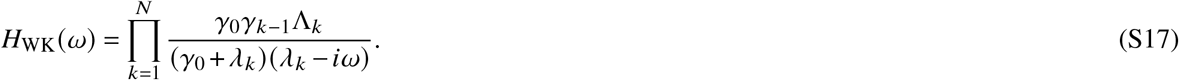

#### 1.4 Calculating the optimal error *E*_WK_

To calculate *E*_WK_ from Eq. (S7), we first take the inverse Fourier transform of *H*_WK_(*ω*) from Eq. (S17), which gives a sum of exponentials in the time domain,

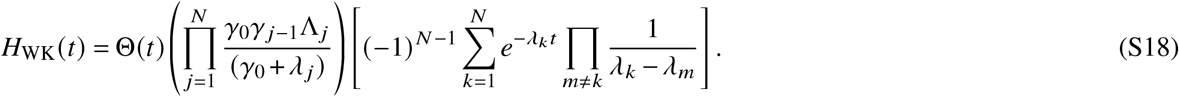

Using the fact that 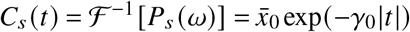, we can evaluate the integral in Eq. (S7) to find

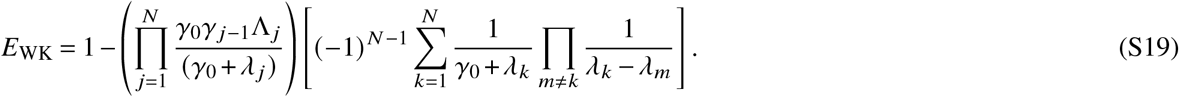

Reversing the partial fraction decomposition,

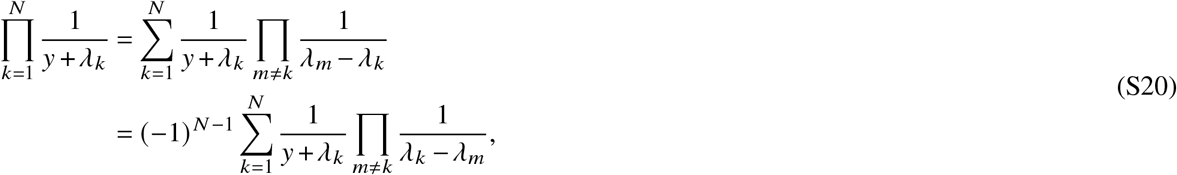

with *y* = *γ*_0_, the error reduces to the value in main text Eq. (13):

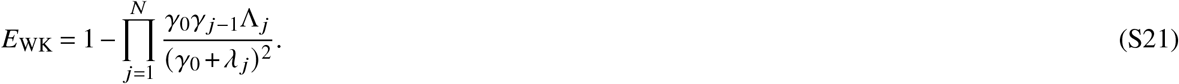

#### 1.5 Conditions under which the system can achieve WK optimality

In order for the system to attain *E* = *E*_WK_, the parameters must be tuned such that *H*(*ω*) ∝ *H*_WK_(*ω*), where *H*(*ω*) and *H*_opt_(*ω*) are given by Eqs. (S5) and (S17) respectively. Comparing the two functions, we see that they are proportional to one another when *λ*_*j*_ = *γ*_*j*_ for all *j* = 1, … , *N*. Satisfying this condition actually requires a certain relationship between the different per-capita deactivation rates *γ*_*j*_ and the Λ_*j*_ parameters.

To see this, let us first denote *B*_*N*_(*λ*) as the polynomial from Eq. (S10) for a particular value of *N*. The explicit forms of the polynomials for the first few values of *N* are as follows:

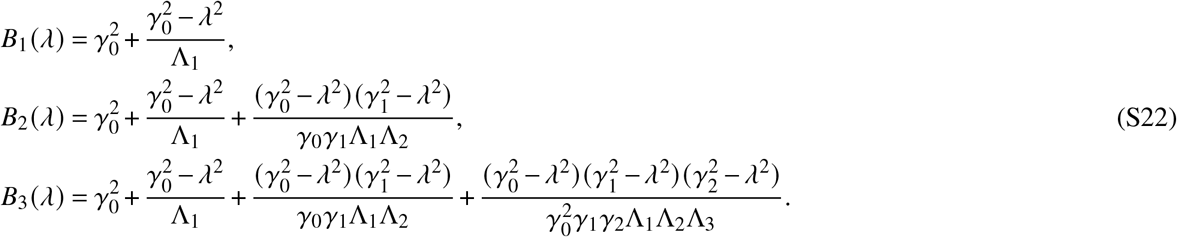

Consider the *N* = 1 system. There is one root *λ*_1_ with a positive real part, and we set it to *λ*_1_ = *γ*_1_ to satisfy the condition. This requires that *B*_1_ (*γ*_1_) = 0, which occurs when 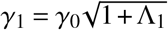. Interestingly, this same value of *γ*_1_ will also be a root for all higher polynomials *N* > 1. Because the additional terms in the higher polynomials all contain a 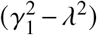 factor, we see that *B*_*N*_ (*γ*_1_) = *B*_1_ (*γ*_1_) = 0 for *N* > 1.

Thus *B*_2_ (*λ*) has one root 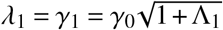 that we have already found, and a new root *λ*_2_ = *γ*_2_ whose value we need to determine. This will be true iteratively at every higher value of *N*: the first *N* − 1 roots *λ*_j_ = *γ*_*j*_, *j* = 1, … , *N* − 1, will be the same roots as for *B*_*N* − 1_(*λ*), and there will one new root *λ*_*N*_ = *γ*_*N*_. This follows from the structure of the *B*_*N*_ (*λ*) polynomials, where

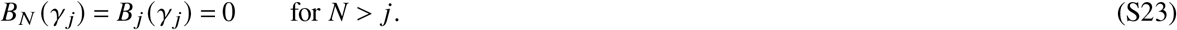

We can find all the higher roots by induction. Let us assume that we have already found the values of *λ*_*j*_ = *γ*_*j*_ for *j* = 1, … , *N* – 1 and are interested in finding *λ*_*N*_ = *γ*_*N*_. The known roots allow us to completely factor *B*_*N* - 1_(*λ*), and from the definition of the polynomials in Eq. (S10) that factorization has to take the form:

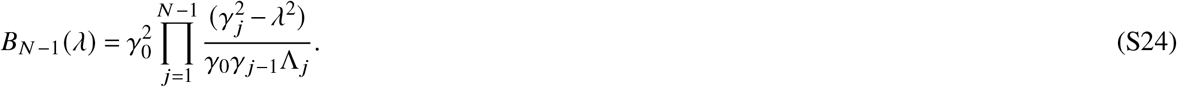

Note that we know the overall prefactor in the factorization above from the prefactor of the highest power *λ*^2(*N*−1)^ in the definition of *B*_*N*−1_ (*λ*). Turning to *B*_*N*_ (*λ*), we can write this polynomial as *B*_*N*−1_ (*λ*) plus an added term,

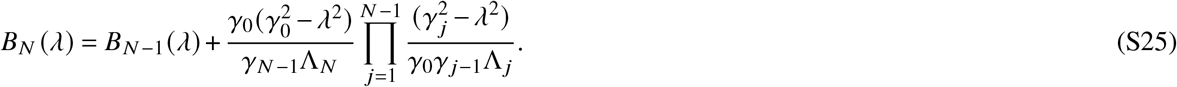

Comparing Eq. (S25) to Eq. (S24), we see that

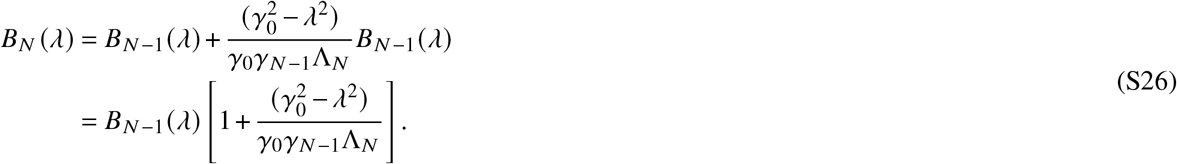

Setting the factor in the brackets to zero allows us to find the new root *λ*_*N*_ = *γ*_*N*_ in terms of the previous root *γ*_*N*−1_,

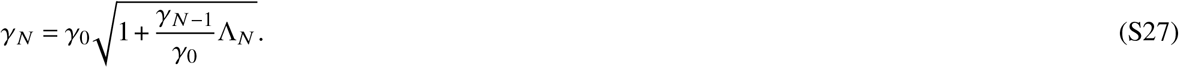

Starting from the known value of 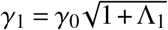, we can iteratively use Eq. (S27) to find all the higher roots. The solutions are the nested radical forms shown in main text Eq. (17),

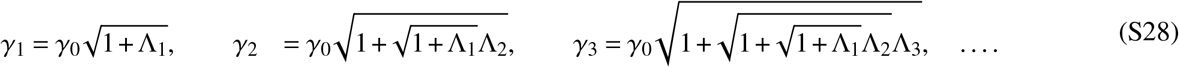

When these conditions are satisfied, the expression for *E*_WK_ simplifies to the form in main text Eq. (18),

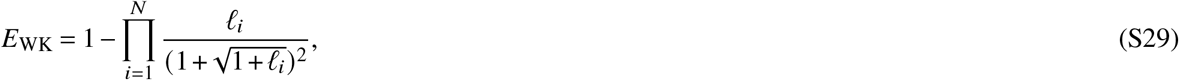

where *l*_*i*_ = *γ*_*i*−1_/Λ_*i*_/*γ*_0_.

### 2 Deriving the WK optimal filter results for the multi-level cascade with feedback

#### 2.1 Mapping the system onto a noise filter, finding the WK filter function and bound

The feedback derivation starts with main text Eq. (9), but with the *ϕ*_1_ term present:

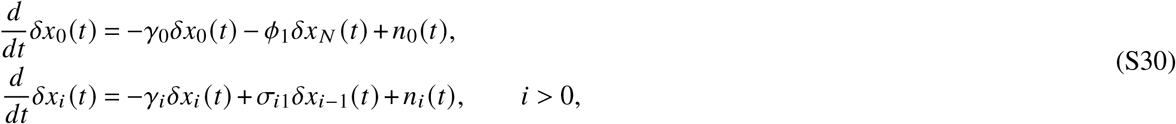

The noise filter mapping is qualitatively different from the no feedback case, taking the form of main text Eq. (19),

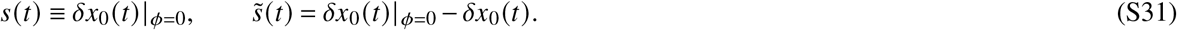

We know the *δx*_0_ (*t*)| _*ϕ*=0_ solution in Fourier space already, having calculated it in Eq. (S3),

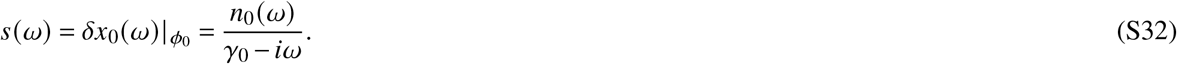

We can manipulate the Fourier space counterpart of Eq. (S30) to relate 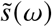 to *s*(*ω*) through a noise filter equation,

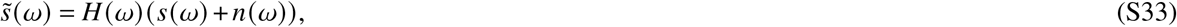

where

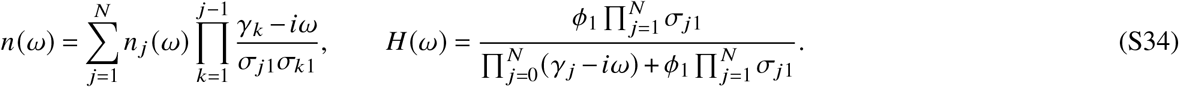

Comparing to Eq. (S5), we see that *s*(*ω*) and *n*(*ω*) in this mapping are exactly the same as in the no feedback case. Hence *P*_*s*_(*ω*) and *P*_*n*_(*ω*) are the same, which means the calculation of *H*_WK_ and *E*_WK_ is unchanged. The result for *E*_WK_ in Eq. (S21) serves as a lower bound for the error *ϵ*.

#### 2.2 Conditions under which the system can achieve WK optimality

Comparing *H*(*ω*) from Eq. (S34) and *H*_WK_(*ω*) from Eq. (S17), one sees that achieving *H*(*ω*) = *H*_WK_(*ω*), and hence *ϵ* = *E*_WK_, is non-trivial. However there is one scenario where this can be approximately fulfilled. We will show that in a certain limit the *N*-level feedback system effectively behaves like an *N* = 1 level system with an effective Λ_1_ parameter. Note that the *N* = 1 version of *P*_*n*_(*ω*) from Eq. (S8b) looks like:

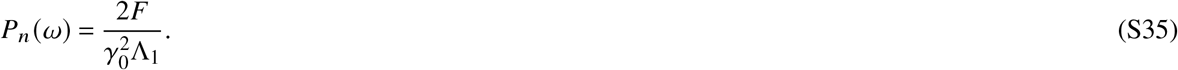

Let us now consider an *N*-level system where *γ*_*j*_ ≫ *γ*_0_ for *j* > 0. The main frequency scale in the system is set by the input signal, which has characteristic frequency *γ*_0_, so typical frequencies *ω* that are relevant to the system behavior all share the property that *ω* ≪ *γ*_*j*_ for *j* > 0. If we use this simplification in Eq. (S8b), the noise power spectrum can be approximated as:

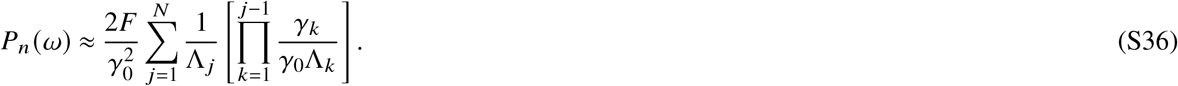

Comparing Eq. (S35) to Eq. (S36), we note that the multi-stage noise power spectrum is approximately the same form as for an *N* = 1 system, except with Λ_1_ replaced by an effective parameter Λ_eff_ given by:

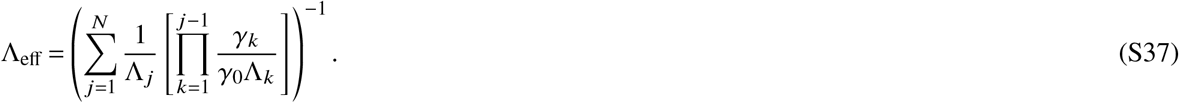

For the special case where the production functions *R*_*j*_(*x*_j-1_) = *σ*_*j*1_*x*_*j*-1_, and hence 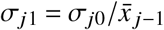 for *j* > 0, the expression for Λ_eff_ simplifies to the result shown in main text Eq. (22):

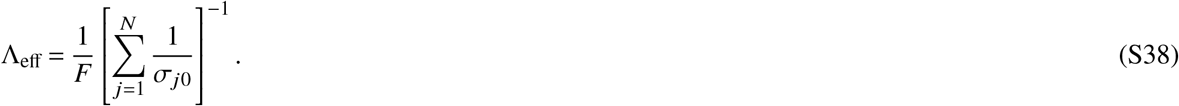

The corresponding *N* = 1 optimal filter *H*_WK_(*ω*) from Eq. (S17), with Λ_eff_ instead of Λ_1_, can be expressed as:

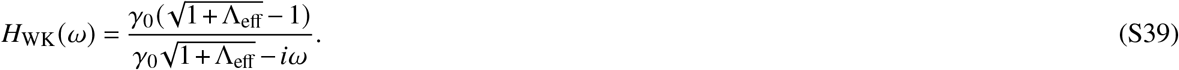

Here we have used the fact that 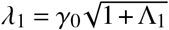 is the root for *B*_1_ (*λ*) from Eq. (S22), and substituted in Λ_eff_.

Let us now write *H*(*ω*) from Eq. (S34) using the approximation *ω* ≪ *γ*_*j*_ for *j* > 0,

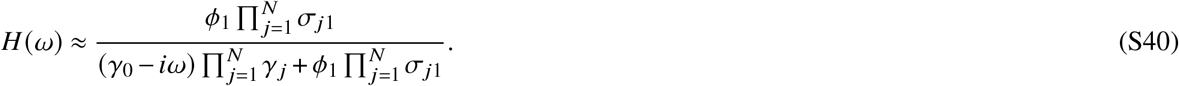

We can thus approximately have *H*(*ω*) ≈ *H*_WK_(*ω*) from Eq. (S39) when the feedback strength is tuned to the value from main text Eq. (21),

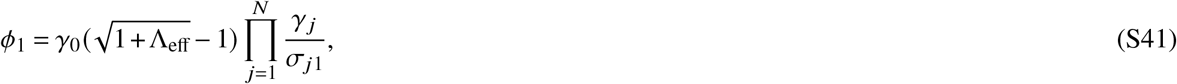

which then ensures that *ϵ* ≈ *E*_WK_, with the latter having the *N* = 1 form,

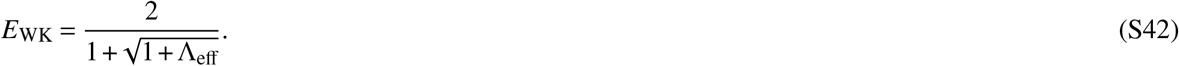

### 3 Exact error calculation in the nonlinear cascade without feedback

This section fills in the details of the calculation that transforms main text Eq. (37), a relation for the generating function 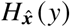 and its derivatives 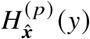, into the recursion relation of main text Eq. (49). The ultimate goal is to use the recursion relation to find the coefficients 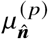 in order to evaluate the exact error *E* given by main text Eq. (48):

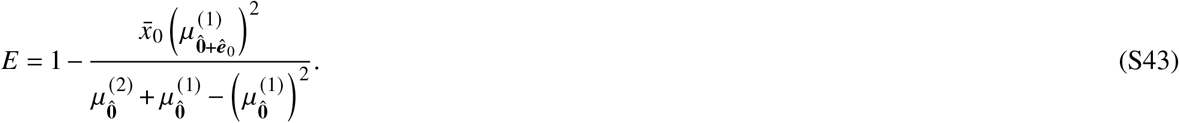

Recall the expansions defined in the main text for all the quantities of interest:

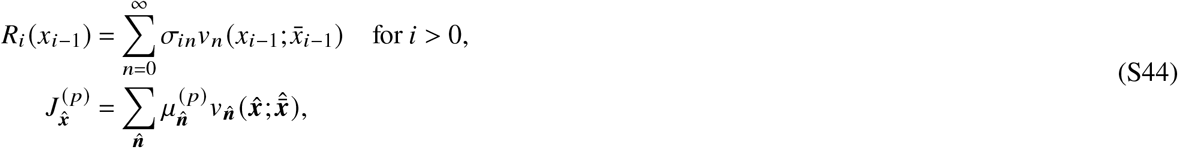

where

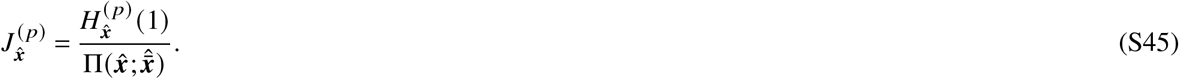

Here we use the multi-dimensional versions of the Poisson distributions and Poisson-Charlier polynomials,

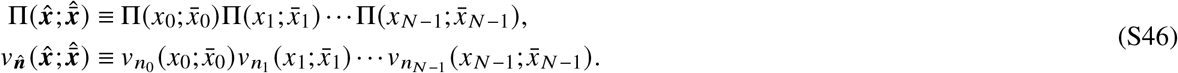

More details on the Poisson-Charlier polynomials can be found in the next section of the SI, which provides a brief guide to their most useful properties.

Since we know the production functions *R*_*i*_ (*x*_*i*−1_) for our system of interest, we can easily find the coefficients *σ*_*in*_ in Eq. (S44), using main text Eq. (42). To derive the coefficients 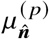 we start with the relation in main text Eq. (37):

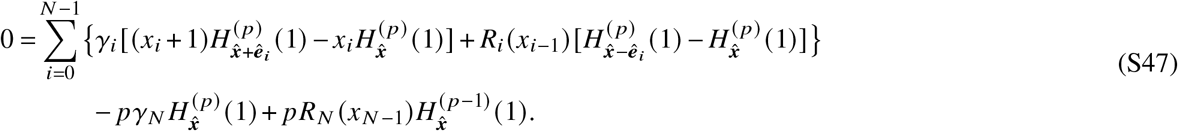

Using Eq. (S45) and the fact that Poisson distributions satisfy 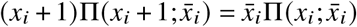, we can rewrite Eq. (S47) in terms of the 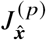 functions:

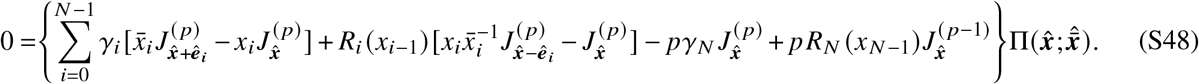

Let us introduce one more expansion, for products of the *R*_*i*_ (*x*_*i*−1_) and 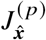 functions,

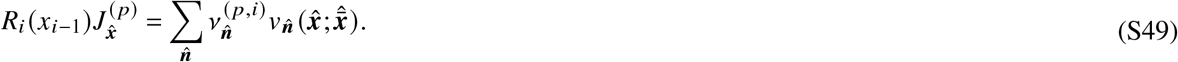

Because *R*_*i*_ (*x*_*i*−1_) and 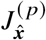 have their own individual expansions in terms of the Poisson-Charlier polynomials, defined by Eq. (S44), the coefficients 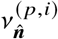 are entirely determined by the coefficients *σ*_*in*_ and 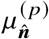 of the individual expansions. This relation, a property of the Poisson-Charlier polynomials, is explained in more detail in SI Sec. 4.5. It takes the form:

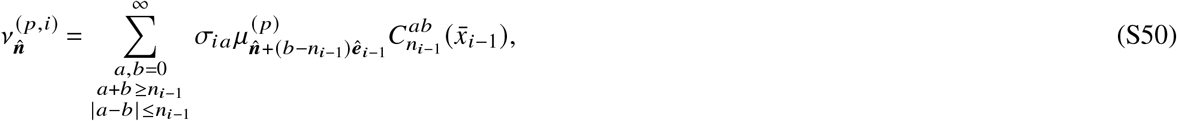

where 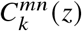 are polynomi defined in Eqs. (S66)-(S67).

Let us define 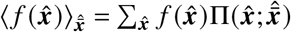 as the average of a function 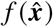 with respect to 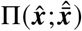. Using the recursion relationships for Poisson-Charlier polynomials shown in Eq. (S64), one can prove the following useful identities:

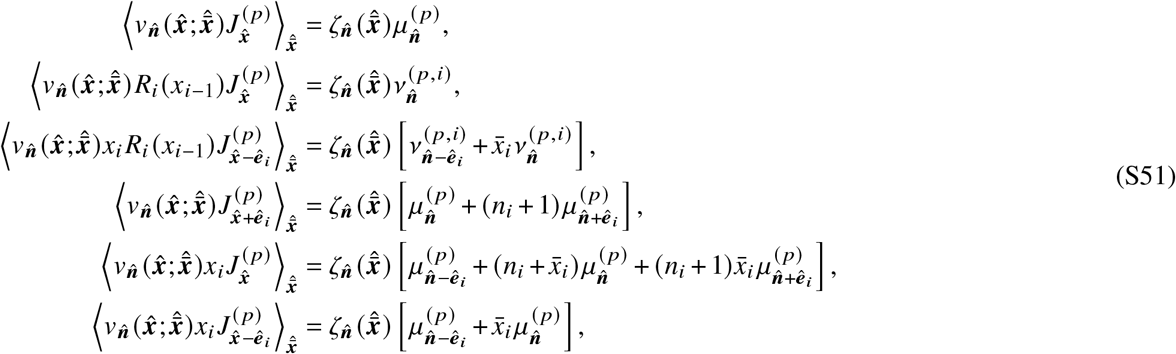

where 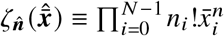. By multiplying Eq. (S48) by 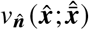 and summing over 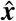, we can use the above averages to obtain the following relation:

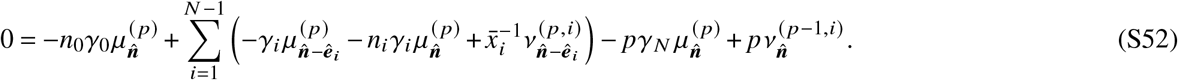

We can rearrange this obtain the recursion relation in main text Eq. (49),

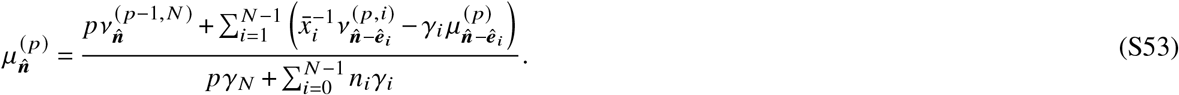

This relation, together with 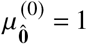 which we know from the normalization property 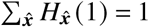, is sufficient for us to calculate any coefficient 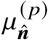 of interest.

### 4 Properties of the Poisson-Charlier polynomials

#### 4.1 Definition of the polynomials

In this section, we summarize some properties of the polynomials 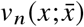 used in our analytical expansion approach for calculating moments of master equations. These are variants of Poisson-Charlier (PC) polynomials^4,5^, 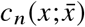, related by a trivial factor to the standard PC definition:

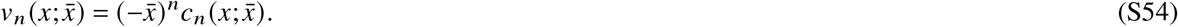

The *n*th function 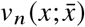 is a polynomial in *x* of degree *n*, depending on the parameter 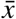. It is defined as follows:

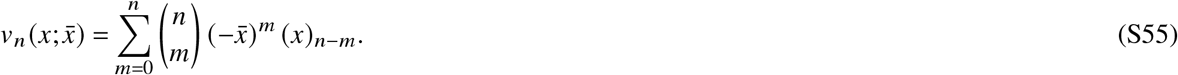

Here 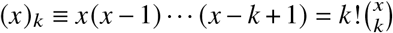 are given by: is the *k*th falling factorial of *x*, with (*x*)_0_ ≡ 1. The first few polynomials

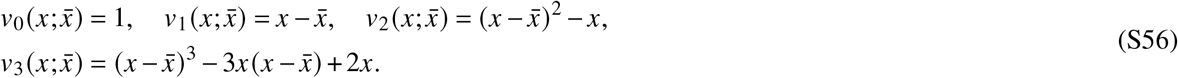

These 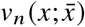 appear in a variety of master equation expansion approaches, for example the spectral method of Refs. 6, 7. In fact, 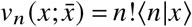, where ⟨*n|x*⟩ is the mixed product defined in Eq. A8 of Ref. 6 (with 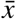 substituted for the rate parameter *g*).

#### 4.2 Orthogonality with respect to the Poisson distribution

One of the convenient properties of these polynomials is that they have simple averages with respect to the Poisson distribution,

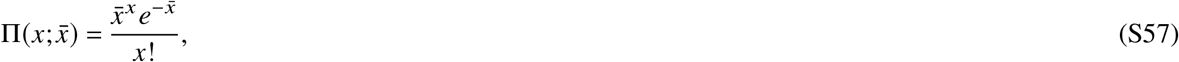

where *x* is a non-negative integer, and 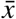 is the parameter that defines the mean of the distribution, so that 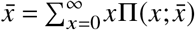. Let us denote the average of a function *f x* with respect to the Poisson distribution 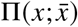 in following way:

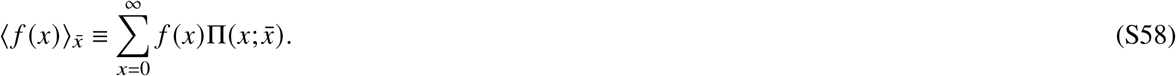

Then the polynomials of Eq. (S55) satisfy the following orthogonality relationship^8, 9^:

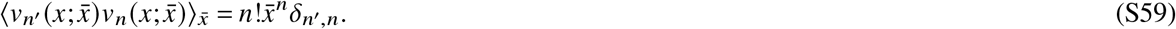

Since 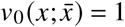, a special case of Eq. (S59) when *n*′ = 0 gives an expression for the mean:

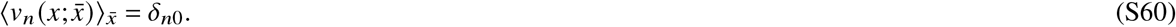

#### 4.3 Using the polynomials as a basis for function expansions

The polynomials form a basis in which one can expand arbitrary functions of populations *f*(*x*),

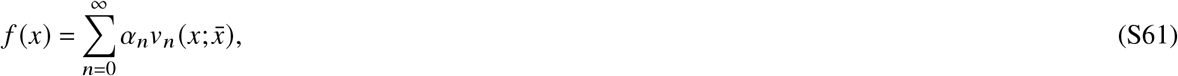

for some coefficients *α*_*n*_. To calculate the *m*th coefficient *α*_*m*_, we multiply both sides of Eq. (S61) by 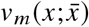 take the average with respect to 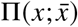:

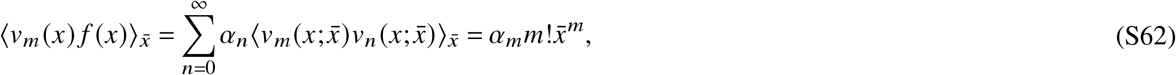

where we have used the orthogonality relation Eq. (S59). Thus *α*_*m*_ is given by:

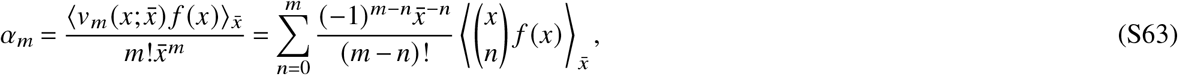

where we have plugged in the definition of 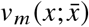 from Eq. (S55). For the kinds of functions we ordinarily encounter in working with master equations, the coefficients *α*_*m*_ rapidly decay with *m*, so in practice we can often form an excellent approximation by just keeping the first few (*n* ≤ 5) terms in the expansion of Eq. (S61)^9^.

#### 4.4 Recursion relationships

The polynomials satisfy the following recursion relationships, as can be easily verified from their definition in Eq. (S55):

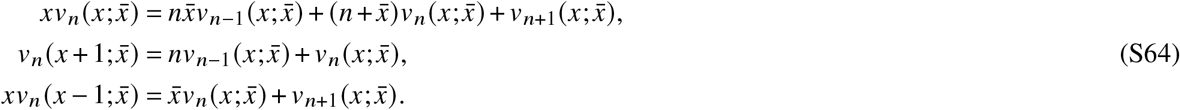

#### 4.5 Expanding the product of polynomials

The final property that comes in useful in calculations is that the product of two polynomials 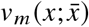 and 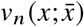 can be itself expanded in a linear combination of polynomials in the following form:

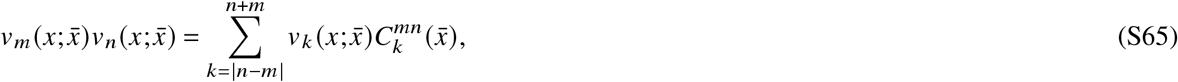

where the coefficients 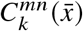 are polynomials in 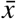 given by:

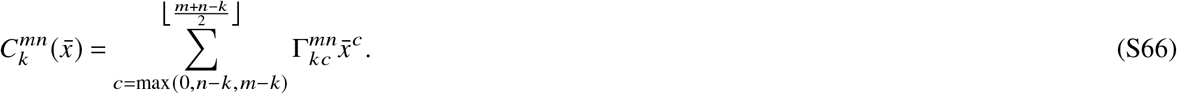

Here, the sum starts at the largest of the three values 0, *n*–*k*, and *m*–*k*, and ⌊*z*⌋ denotes the largest integer less or equal to *z*. The quantity 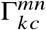 is defined as:

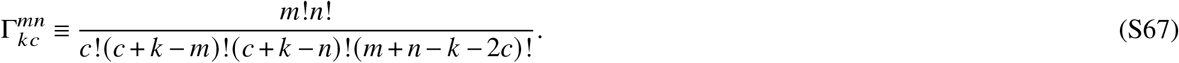

Thus for example if one had two functions *f*(*x*) and *g*(*x*) with individual expansions,

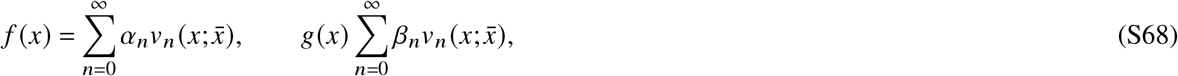

then the product can be expanded as

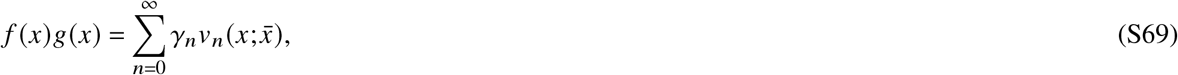

with coefficients given by

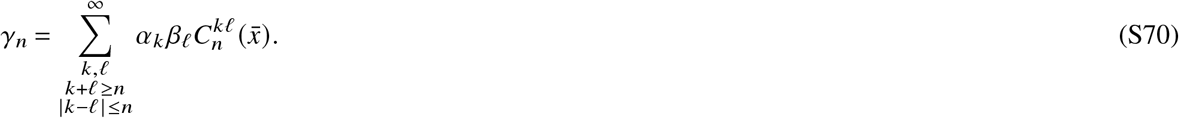

